# Nutrient retention by predators undermines predator coexistence on one prey

**DOI:** 10.1101/535195

**Authors:** Toni Klauschies, Ursula Gaedke

## Abstract

Contemporary theory of predator coexistence through relative non-linearity in their functional responses strongly relies on the Rosenzweig-MacArthur equations (1963) in which the (autotrophic) prey exhibits logistic growth in the absence of the predators. This implies that the prey is limited by a resource such as light or space, which availability is independent of the predators. This assumption does not hold under nutrient limitation where both prey and predators bind resources such as nitrogen and phosphorus in their biomass. Furthermore, the prey’s resource uptake-rate is assumed to be linear and the predator-prey system is considered to be closed. All these assumptions are unrealistic for many natural systems. Here, we show that nutrient retention in predator biomass strongly hampers the coexistence of two predators on one prey because it stabilizes the dynamics. In contrast, a non-linear resource uptake rate of the prey slightly promotes predator coexistence. Our study highlights that predator coexistence does not only depend on differences in the curvature of their functional responses but also on the type of resource constraining the growth of their prey. This has far-reaching consequences for the relative importance of fluctuation-dependent and -independent mechanisms of species coexistence in natural systems where autotrophs experience light or nutrient limitation.

## 1. Introduction

Since the seminal work of Lotka (1925), Volterra (1928), Gause (1932) and Hutchinson (1961) ecologists are wondering about the mechanisms that underlie the astonishing diversity observed in natural systems. Contemporary theory distinguishes two broad categories of mechanisms that promote species coexistence. Fluctuation-independent mechanisms such as resource partitioning or specialist predation may allow different species to coexist under constant environmental conditions (Chesson 2000; Chase and Leibold 2003). This contrasts with fluctuation-dependent mechanisms that rely on temporal variation in environmental conditions (Abrams and Holt 2002; Chesson 2003). For example, species may be able to coexist due to species-specific differences in their demographic responses to exogenously driven fluctuations in the abiotic environment (Chesson and Warner 1981; Ellner et al. 2016) or because of endogenously generated oscillations in species abundances (Armstrong and McGehee 1980; Yamamichi et al. 2011) or traits (Abrams 2006; Klauschies et al. 2016). In this study, we investigate how the type of resource limiting prey growth influences the coexistence of two predators on one fluctuating prey through relative non-linearity in their functional responses, as it represents an important food web module in more complex and diverse communities.

Previous work demonstrated that two predators can coexist on one prey for a broad range of parameters when the system exhibits pronounced predator-prey oscillations and when the predators show substantial differences in the curvature of their functional responses (Koch 1974; Hsu et al. 1978; Armstrong and McGehee 1980; Abrams and Holt 2002). The predator with the more linear functional response is the better competitor when the system exhibits strong predator-prey oscillations but promotes less variation in abundances when it reaches high densities. In contrast, the predator with the more non-linear functional response is competitive superior at steady state dynamics while favoring larger fluctuations in abundances when it becomes dominant (Abrams 2004; Mittelbach 2012; Xiao and Fussmann 2013). Hence, the dominant predator alters the environment in a way that is beneficial for the rare predator, allowing its regrowth from low densities (Figure 1A). This negative frequency-dependent selection may enable stable coexistence of the two different predators: in line with the gleaner-opportunist trade-off (Grover 1997; Litchman and Klausmeier 2001), one predator (the gleaner) exhibits a higher net-growth growth rate at lower prey densities whereas the other predator (the opportunist) benefits from a higher net-growth rate at higher prey densities.

**Figure 1:**
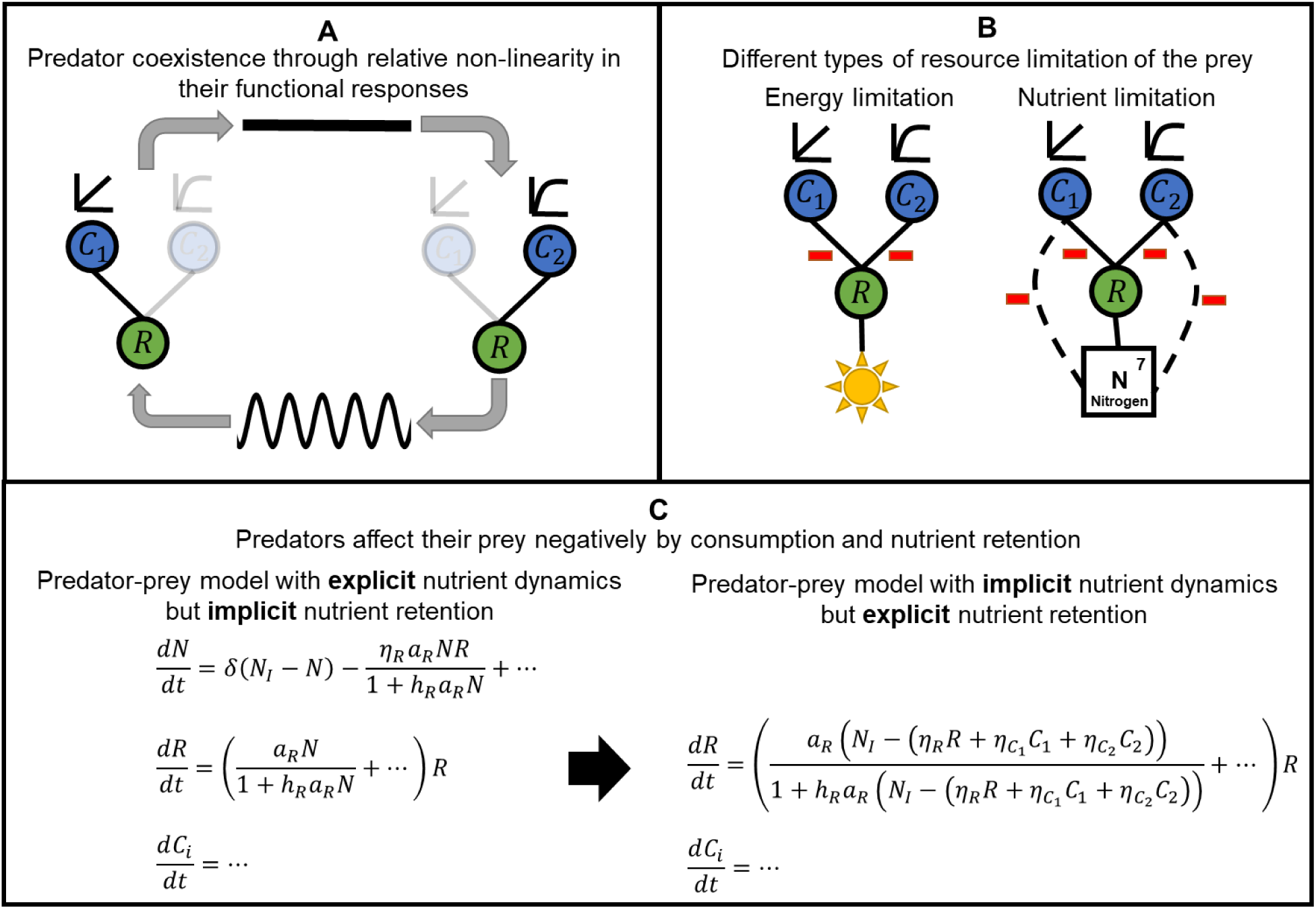
Predator coexistence through relative non-linearity in their functional responses and associated fluctuations in species abundances. (A) Negative frequency-dependent selection: the dominant predator (in-transparent) alters the environment in a way that is beneficial for the rare predator (transparent). A dominance of the predator with a type I functional response promotes stasis in the predator-prey dynamics although it is the better competitor under fluctuating environmental conditions. In contrast, a dominance of the predator with a type II functional response promotes predator-prey oscillations while being competitive superior under constant environmental conditions. (B) The type of resource limitation influences predator-prey interactions beyond consumptive effects of the predators on the prey. Predators mainly affect the prey by consumption under light or energy limitation whereas under nutrient limitation of the prey, predators also impact the prey negatively by nutrient retention. (C) The predator-prey model defined by eq. [9] in the main text can be derived from a generic predator-prey model that explicitly accounts for the replenishment, uptake, excretion and recycling of nutrients (for details see methods).

Most theoretical investigations in this field rest on the classical equations of Rosenzweig and MacArthur (1963) where the prey grows logistically in the absence of the two predators (e.g. Hsu et al. 1978; Xiao and Fussmann 2013). This implies three very strong assumptions about the type, the uptake and the replenishment of the resource that is limiting the growth of the (autotrophic) prey, which may or may not hold in nature.

First of all, logistic growth entails that the predators will not directly affect the amount of resources freely available for prey uptake (DeAngelis 1992). This is reasonable for resources such as light or space, which are hardly influenced by predators. However, when mineral nutrients such as nitrogen or phosphorus are limiting autotrophic or bacterial growth, the resource is often substantially stored within the predator and prey biomass. As a result, the predators affect the prey not only by consumption but also by nutrient retention (Figure 1B; DeAngelis 1992). Given that herbivores have much higher nutrient to carbon ratios (N/C ratios) than nutrient depleted plants (Sterner and Elser 2002; Andersen 2004; Hessen et al. 2013), nutrient retention by predators may substantially influence prey growth even when the total biomass of the predators is relatively low (Gaedke et al. 2002).

Second, under logistic growth, the resource uptake rate of the prey is assumed to be linear (Kooi et al. 1998a). This assumption lacks empirical evidence (Eppley et al. 1969; Sibly and Hone 2002) and is not supported by mechanistic considerations (Dugdale 1967; Droop 2003). Rather, in line with classical enzyme kinetics (Michaelis and Menten 1913) and functional response theory (Holling 1961), it typically depends non-linearly on the external resource concentration. A linear uptake rate is only to be expected at very low resource concentrations, e.g. when the prey is strongly bottom-up controlled whereas, at higher resource concentrations, it is likely to be saturating.

Finally, the Rosenzweig-MacArthur predator-prey model basically describes a closed autochthonous system with a given constant amount of resources, the prey’s carrying capacity K. In this case, nutrients are thought to be internally replenished. However, most natural systems receive allochthonous nutrient inputs and experience losses of dissolved and organically bound nutrients, i.e. they are more realistically described as flow-through systems. Hence, we investigate to what extent previous results on the coexistence of two predators sharing a single prey depend on the above mentioned simplifying assumptions. We show that coexistence is most likely in autochthonous systems where the predators do not interfere with the resource availability of the prey, e.g. under light or space limitation, and when the preys’ uptake rate for the resource is strongly non-linear. Therefore, predator coexistence does not only depend on differences in the non-linearity of their functional responses but also on the type of resource constraining the growth of their prey. This is crucial for our understanding of species coexistence in natural systems because we can expect that fluctuation-dependent mechanisms of species coexistence are more relevant in light or space than nutrient limited systems.

## 2. Methods

We derive a generic predator-prey model that integrates and extends previous studies investigating predator coexistence on one prey through differences in their functional responses and endogenously generated fluctuations in species abundances under batch (Koch 1974; Armstrong and McGehee 1980; Hsu et al. 1978; Abrams and Holt 2002; Xiao and Fussmann 2013), or chemostat conditions (Butler and Waltman 1981; Butler et al. 1983; Keener 1985).

### 2.1. Description of the predator-prey model

First, we consider a universal predator-prey system that comprises two different predators and one prey and explicitly accounts for the replenishment, uptake, excretion and recycling of inorganic mineral nutrients such as nitrogen or phosphorus in a flow-through (chemostat) system. Hence, the concentration of dissolved inorganic nutrients *N* and the biomasses of the prey *R* and the two predators *C*_1_ and *C*_2_ are changing over time according to the following set of equations:

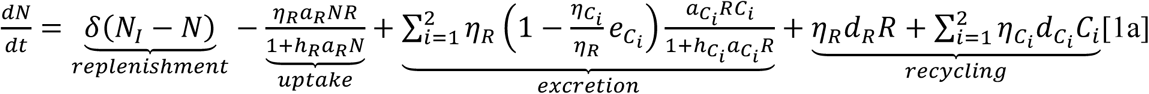

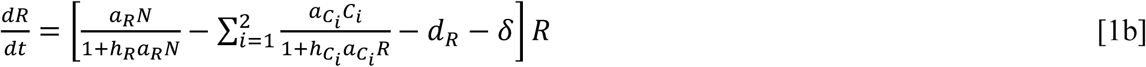

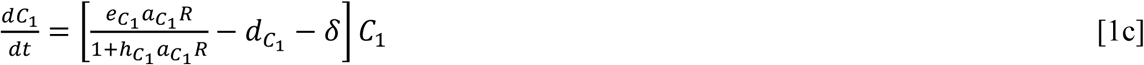

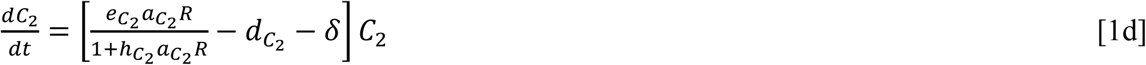

The rate of change of *N*, i.e. *dN*/*dt*, depends on the dilution rate *δ*, the concentrations of dissolved inorganic nutrients in the inflowing medium *N*_*I*_ and in the chemostat *N*, the nutrient-uptake of the prey, the excretion of nutrients by the predators and the recycling of nutrients contained in dead predators and prey. Excretion and recycling depend on the nutrient to carbon ratios 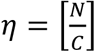 of the different species, which we assumed to be constant. When the nutrient to carbon ratios are equal for the different species, the excretion term reduces to 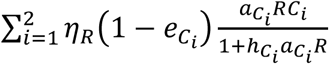 (cf. DeAngelis 1992). However, when they are different, our general formulation is more accurate. For example, it is realistic to assume that the predators contain more nutrients per unit carbon than the autotrophic prey, which reduces the nutrients excreted by the predators.

Furthermore, nutrient-uptake of the prey follows a type II functional response, i.e. Michaelis-Menten-Kinetik (Michaelis & Menten 1913), with uptake rate *a*_*R*_ and handling time *h*_*R*_ (Holling, 1961). Inorganic nutrients are converted into prey biomass according to its specific nutrient to carbon ratio *η*_*R*_. The two predators also feed on the prey with a type II functional response with species-specific attack rates 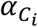 and handling times 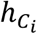. The ingested prey biomass is converted into predator biomass with species-specific transfer efficiencies 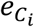. To reflect most natural conditions, predators are assumed to be energy- and thus, carbon-limited. Hence, the transfer efficiencies of the predators ensure that the stoichiometry of the prey meets the nutritional demands of the predators, i.e.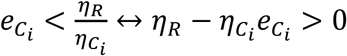. Hence, the food quality of the prey, i.e.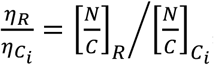, is sufficiently high to support the energetically feasible biomass production of the predators in units of carbon. Therefore, the loss of energy through respiration or low assimilation efficiencies of carbon prevents further growth of the predator biomasses so that excretion may substantially supply dissolved inorganic nutrients.

In addition, all nutrients and organisms are washed out from the system with the dilution rate *δ*, constituting a shared loss rate for all species. Finally, we also accounted for senility or basal mortality of the different species by incorporating the species-specific death rates *d*_*R*_ and 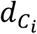 for the prey and the two predators, respectively. Accounting for nutrient excretion by the predators and the recycling of nutrients from dead organisms ensures that the predator-prey model described by eq. [1] is mass-balanced.

### 2.2. Model simplification

In order to facilitate model analysis we reduced the number of state variables and parameters of our general predator-prey model.

#### 2.2.1. Reducing the number of state variables

The predator-prey model defined by eq. [1] contains a conserved quantity, which allows us to reduce its dimension. Following the methodology used by Armstrong and McGehee (1980), Jones and Ellner (2007), Scranton and Vasseur (2016) and O’Dwyer (2018), we first define the total amount of nutrients, i.e. the sum of dissolved inorganic and organically bound nutrients, in our system as follows: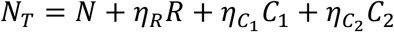. Consequently, the rate of change of *N*_*T*_ is given by:

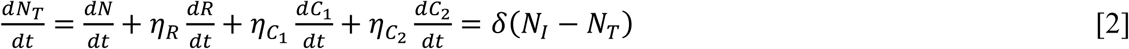

which can be solved analytically by substitution with 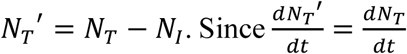 we get:

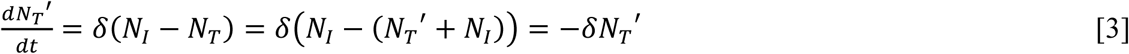

This equation represents exponential growth, which has the solution:

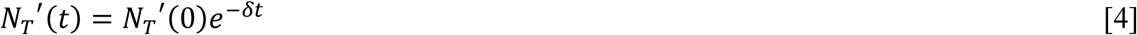

Using the backwards substitution terms *N*_*T*_(*t*) = *N*_*T*_′(*t*) + *N*_*I*_ and *N*_*T*_′(0) = *N*_*T*_(0) − *N*_*I*_ we can now express *N*_*T*_(*t*) as:

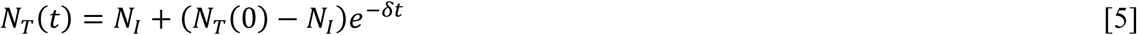

In the limit as *t* → ∞ the system reaches its asymptotically stable equilibrium: *N*_*T*_(*t*) → *N*_*I*_. Therefore, we can write a steady-state approximation for *N*_*T*_:

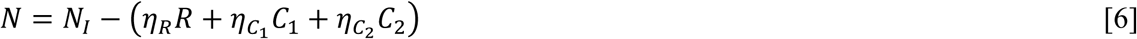

Substituting eq. [6] into eq. [1] results into the dimensionally reduced predator-prey model:

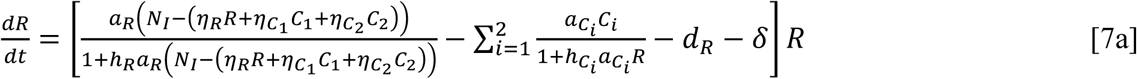

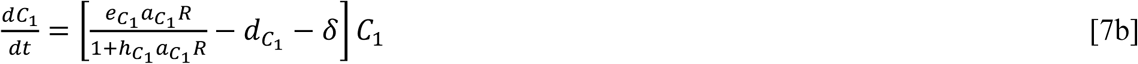

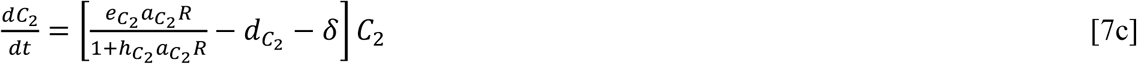

The concentration of dissolved inorganic nutrients which are freely available to the prey, is given by 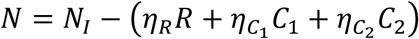 where *N*_*I*_ denotes the total amount of nutrients in the system. The term 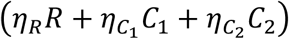 quantifies the amount of nutrients contained in the species biomasses, which are not available for prey growth. Hence, predators not only feed on the prey but also diminish the nutrients available for their own prey by nutrient retention (Figure 1B, C). This effectively reduces the carrying capacity of their prey. The overall relevance of this effect will generally depend on the nutrient to carbon ratios of the different species and to what extent mineral nutrients are limiting prey growth. The negative effect of nutrient retention may surpass that of predation when the predators possess low feeding rates and comparably high nutrient to carbon ratios. In contrast, when light or space is limiting prey growth, the two predators do not influence the amount of the limiting resource because they do not contribute to self-shading or competition for space.

In line with Lotka-Volterra competition models (cf. MacArthur and Levins 1967; Schoener 1974) the terms describing nutrient retention by the predators in the prey equation are very akin to a phenomenological description of resource competition. Hence, the nutrient to carbon ratios of the prey (*η*_*R*_) and the predators 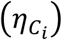 may also be interpreted as species-specific competition coefficients for the total amount of resources.

The two predator-prey systems defined by eq. [1] and eq. [7] are equivalent when the dilution rate is larger than zero. When choosing the same initial conditions, both models predict the same transients and equilibrium dynamics. The only restriction is that the initial amount of inorganic nutrients of eq. [1] has to be chosen to be the difference between the total amount of nutrients in the predator-prey system and the initial amount of organically bound nutrients, i.e.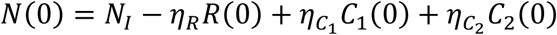. Otherwise the two models may show differences in the transients of the their population dynamics. However, differences in the equilibrium dynamics of the two different models can only occur when the dilution rate is set to zero, because in this case, the total amount of nutrients in the model defined by eq. [1] depends on the initial amount of organically bound nutrients and dissolved inorganic nutrients in the system.

#### 2.2.2. Reducing the number of parameters by non-dimensionalization

We further simplified the analysis by rescaling the time, state variables and parameters of our predator-prey model as follows: 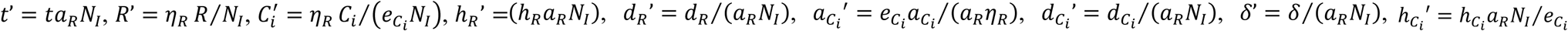 and 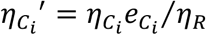. Substituting the rescaled state variables and effective parameters into eq. [7] and dropping the primes, we obtain the following non-dimensionalized predator-prey model:

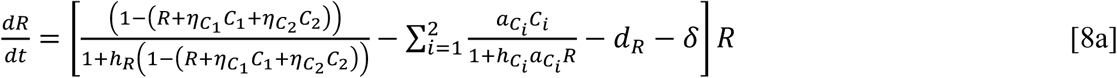

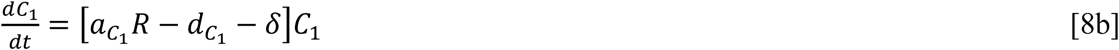

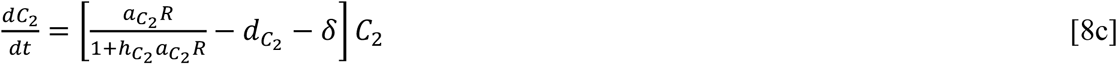

Hence, the overall population dynamics of the different species depend on the curvature of the functional responses and the nutrient to carbon ratios of the two different predators, the non-linearity of the nutrient uptake rate of the resource, the species-specific death rates and the shared loss rate for all species. In contrast, the total amount of nutrients in the system, the transfer efficiencies of the two predators, the resource uptake rate of the prey and the nutrient to carbon ratio of the prey can be omitted from further mathematical analysis. Nevertheless, since non-dimensional variables still depend on dimensional variables, any biological analysis should also consider these functional relationships. For example, since 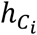′ is inversely proportional to 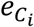, any increase in the efficiency of predator *i* has the same effect as a decrease in 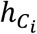.

#### 2.2.3. Simplifying assumptions

Previous studies have shown that two predators are most likely to coexist on a single prey through endogenously generated predator-prey oscillations when they exhibit substantial differences in their functional responses (Hsu et al. 1978; Chesson 2000; Abrams and Holt 2002; Xiao and Fussmann 2013). Hence, we assumed the grazing rate of *C*_1_ to follow a linear, type I, functional response with attack rate 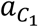 and thus, restricted our analysis to the case where *C*_1_ is not limited by handling the prey, i.e.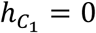. In contrast, the grazing rate of *C*_2_ was assumed to follow a type II functional response with attack rate 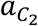 and handling time 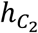 (Armstrong and McGehee 1980; Abrams and Holt 2002).

In addition, the parameter region for which the two different predators may coexist also increases with a higher carrying capacity of the prey, i.e. the equilibrium biomass of the prey in the absence of the two predators (Hsu et al. 1978). This results from the fact that a higher carrying capacity promotes unstable population dynamics, which are necessary for the two predators to coexist due to differences in their functional responses (for details see Appendix A). Given that an increase in *d*_*R*_ effectively reduces the carrying capacity and thereby the likelihood of predator coexistence, we focused our analysis on the case, where the basal prey mortality was negligible compared to the overall losses through predation and dilution, i.e. *d*_*R*_ = 0, which also facilitates comparison with previous results. These modifications simplify our predator-prey model as follows:

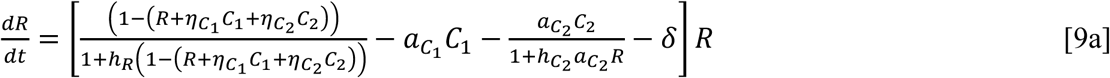

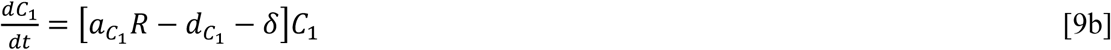

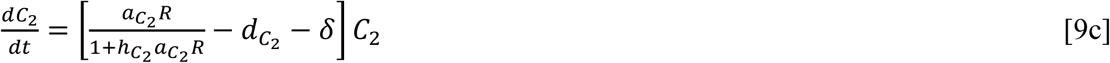

When assuming the absence of nutrient retention by the predators 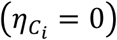, a linear nutrient uptake rate of the prey (*h*_*R*_ = 0), and a closed predator-prey system (batch conditions where *δ* = 0), our system described by eq. [9] reduces to the predator-prey system studied extensively in previous work (e.g. Armstrong and McGehee 1980; Abrams and Holt 2002; Xiao and Fussmann 2013). In this case, the intrinsic growth rate (*r*) and the carrying capacity (*K*) of the prey are given by *r* = *a*_*R*_*N*_*I*_ and *K* = *N*_*I*_. Hence, the classical predator-prey model based on the Rosenzweig-MacArthur equations (Rosenzweig and MacArthur 1963) is a special case of our general model described by eq. [9]. This allows us to directly compare our results to previous findings.

### 2.3. Model analysis

We combined analytical and numerical approaches to achieve a clear and comprehensive understanding of the impact of nutrient retention by the predators and of a non-linear nutrient uptake rate of the prey on the coexistence of two predators on one prey in a batch or a flow-through (chemostat) system.

Predator coexistence on one prey mainly depends on the death rates of the two predators when their functional responses are fixed (Abrams and Holt 2002). We thus evaluated predator coexistence in the two-dimensional 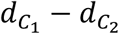 parameter space. In line with Hsu et al. (1978) and Xiao and Fussmann (2013), we first derived the necessary conditions for coexistence that define a range of parameters which may permit coexistence (for details see section 2.3.1 and Appendix A). Within this parameter space, we used numerical simulations to obtain also the sufficient conditions that define the range of parameters enabling coexistence (for details see section 2.3.2). In accordance with previous studies we first investigated the influence of the curvature of the type II functional response of *C*_2_ on predator coexistence assuming logistic growth of the prey in the absence of the predators, thereby setting the baseline for further analysis. Afterwards, we explored the impact of (1) resource retention by the predators 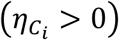, (2) a non-linear resource uptake of the prey (*h*_*R*_ > 0) and (3) a shared loss rate for all species (*δ* > 0) on the coexistence of two predators on one prey (for details see section 2.3.3).

#### 2.3.1. Parameter range of potential species coexistence

In the following we derive necessary conditions defining a region in the 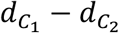 parameter space that may allow predator coexistence. A basic requirement for coexistence is that each predator is able to persist with the prey alone in the absence of the other predator. This prerequisite gives rise to upper limits in the death rates of the two predators as they are given by 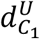 for *C*_1_ and 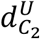 for *C*_2_ (for details see Appendix A.1; cf. Figure 2A – upper limits of the x and y axes).

**Figure 2:**
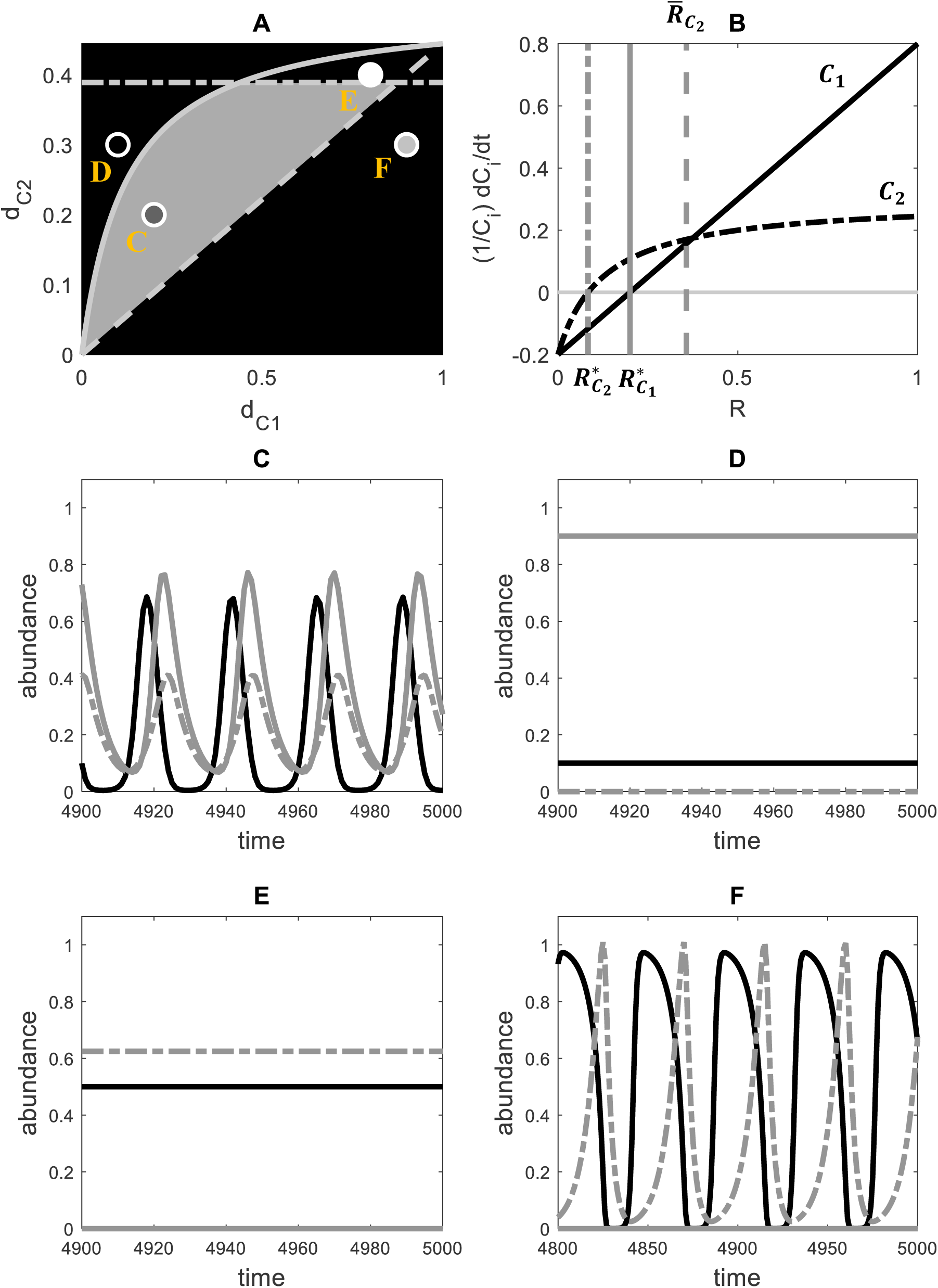
Necessary conditions for the coexistence of two different predators through relative non-linearity in their functional responses and associated fluctuations in the prey abundance R. (A) Parameter range, i.e. death rates 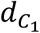 and 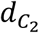, that potentially enables coexistence of predator one, *C*_1_, and predator two, *C*_2_ (P_a_; grey shaded area). The scales of 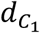 and 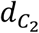 are chosen in accordance with the first necessary condition requiring each predator to persist in the absence of the other one. In line with the second necessary condition *C*_2_ can invade the R − *C*_1_ system when 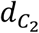 is lower than a threshold value that depends on 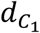 (grey solid line). The third necessary condition predicts the exclusion of *C*_1_ when 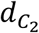falls below a critical value that is also depending on 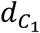 (grey dashed line). Finally, according to the fourth necessary condition, predator coexistence is only possible when the R − *C*_2_ system exhibits predator-prey oscillations. For this, 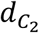 has to be lower than the Hopf-bifurcation point 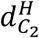 of the R − *C*_2_ system (grey dashed-dotted line). Dots indicate parameter combinations used for the individual simulations shown in (C-F) (for details see methods). (B) Mutual invasibility of the two predators depends on the predators’ per-capita net-growth rates. While *C*_1_ is assumed to exhibit a linear functional response (black solid line), *C*_2_ has a non-linear functional response (black dashed-dotted line). *C*_2_ can invade the R − *C*_1_ system when its minimum prey requirement (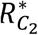, vertical dashed-dotted line) is lower than the minimum prey requirement of *C*_1_ (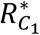, vertical solid line). In contrast, *C*_1_ can invade the R − *C*_2_ system when the abundances of R and *C*_2_ exhibit oscillations and 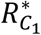 is lower than the temporal average of the corresponding prey density (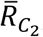, vertical dashed line). (C-F) Dynamics of R (black solid line), *C*_1_ (grey solid line) and *C*_2_ (grey dashed-dotted line) in dependence of the predators’ death rates. For 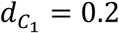 and 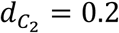 (dark grey dot in (A)), both predators are able to coexist on a limit cycle (C). For 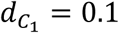and 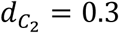 (black dot in (A)), *C*_1_ outcompetes *C*_2_ (D). In contrast, *C*_2_ excludes *C*_1_ for 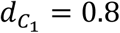 and 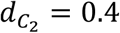 (E, stable equilibrium; white dot in (A)) as well as for 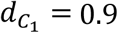 and 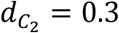 (F, predator-prey oscillations; light grey dot in (A)). All simulations are based on the following parameter values: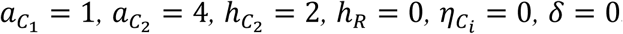.

Furthermore, species are generally viewed to stably coexist when they can mutually invade each other’s resident community (Chesson 2000). Hence, there has to be negative frequency-dependent selection on the different predators, which allows all species to regrow from low densities. The linear functional response of *C*_1_ ensures that the *R* − *C*_1_ system approaches a stable equilibrium in which the equilibrium density of the prey 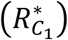 represents the minimum prey requirement of *C*_1_. Therefore, *C*_2_ will be able to invade the *R* − *C*_1_ system when it exhibits a positive per-capita net growth for 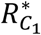. This implies that the minimum prey requirement of 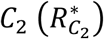 has to be lower than the prey requirement of 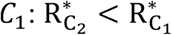 (Appendix A.2; Figure 2A – solid line).

The non-linear functional response of *C*_2_ may destabilize the interior equilibrium of the *R* − *C*_2_ system, provoking predator-prey oscillations. However, due to *C*_1_’s linear functional response, its invasion growth rate for the *R* − *C*_2_ system equals its per-capita net-growth rate for the temporal average of the prey density 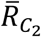. Hence, *C*_1_ can invade the *R* − *C*_2_ system when its per-capita net-growth rate for 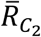 is larger than zero and, thus, when its minimum prey requirement is lower than 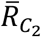, i.e.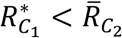. Unfortunately, a general expression for 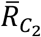 does not exist. We can, however, derive an upper limit for 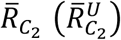 which generally depends on the death rate and the parameters of the functional response of *C*_2_. Hence, *C*_1_ cannot invade the *R* − *C*_2_ system for 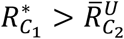 (Appendix A.3; Figure 2A – dashed line).

In line with the reasoning above, the two predators are able to coexist on one prey when the following inequality is satisfied: 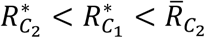 (cf. Figure 2B; Abrams and Holt 2002). Hence, the minimum prey requirement of *C*_2_ has to be lower than the temporal average of the prey density in the *R* − *C*_2_ system, i.e.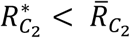. This inequality only holds when the equilibrium value 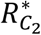 is unstable, i.e., the corresponding predator-prey system exhibits oscillations in the species’ abundances. Hence, the death rate of *C*_2_ has to be smaller than the threshold value 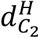, at which the *R* − *C*_2_ system exhibits a Hopf-bifurcation which gives rise to a limit cycle in the corresponding predator-prey dynamics (Figure 2A – dashed-dotted line; for details see Appendix A.4).

As detailed in Appendix B, the differences 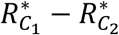 and 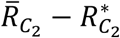 measure the average fitness and niche differences between the two different predators, respectively. Hence, whenever the average fitness differences between the two different predators are smaller than their niche differences, i.e.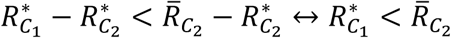, the predators will be able to coexist on one prey under recurrent predator-prey oscillations. Otherwise, one of the two predators will be excluded from the system.

In line with Xiao and Fussmann (2013), the necessary conditions outlined above specify a region in the parameter space in which predator coexistence is possible (cf. Figure 2A – grey shaded area). The absolute size of this parameter region (*P*_*a*_) quantifies the overall strength of relative non-linearity in the predators’ functional responses and associated fluctuations in the prey’s abundance to allow predator coexistence (for details on the quantification of *P*_*a*_ see Appendix A.5). In addition, we calculated a relative size of this parameter region 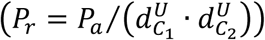 that scales *P*_*a*_ with respect to the upper limits of the predators’ death rates 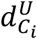. Hence, *P*_*r*_ quantifies the proportion of the parameter space of predator persistence that enables predator coexistence (Figure 2A). Assuming that each predator is able to persist with the prey alone and that the death rates of the different predators are equally likely within the finite two-dimensional 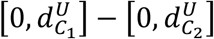 space (uniform distribution), *P*_*r*_ provides a probability that the two predators are able to coexist on one fluctuating prey.

#### 2.3.2 Parameter range of realized species coexistence

Since 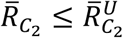, the parameter range of potential species coexistence *P*_*a*_ will generally overestimate the parameter region that actually allows predator coexistence. The magnitude of 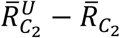 determines to what extent the range of potential coexistence differs from the range of realized coexistence. Hence, we quantified the latter by conducting numerical simulations of our predator-prey model defined by eq. [9].

Following Abrams (2004), we considered predator coexistence to be stable when each predator is able to invade the resident system comprising the other predator and the prey at its long-term equilibrium, combined with persistence of the two predators after invasion. First, we tested for mutual invasibility of the two predators by conducting numerical simulations where either *C*_1_ or *C*_2_ was set to zero at the beginning. The initial density of the resident predator species was set to 0.2 whereas the initial prey density was chosen to equal 0.1. Based on the simulation results of the two different resident systems, we calculated the invasion growth rates, i.e. the time-averaged per-capita net-growth rates, of the two predators according to the following set of equations:

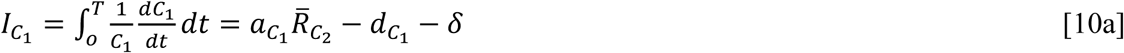

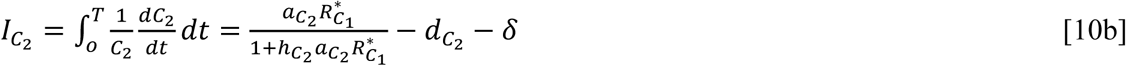

with 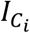 being the invasion growth rate of predator *i* for the corresponding resident system. The overall simulation time *T* was set to 10^4^. The two predators can mutually invade each other when both of them exhibit positive invasion growth rates for the corresponding resident systems, i.e.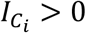. In contrast, if any 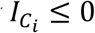, the relevant predator will not be able to recover from low densities, and thus, cannot invade.

We complemented this approach by also testing for persistence of the two predators after invasion, by conducting additional numerical simulations where all species were initially present. Resolving the 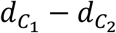 parameter space with a grid of 50 equally spaced values of 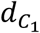 and 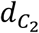, respectively, resulted in a total amount of simulations (*Z*_*total*_) of 50 · 50 = 2500. All simulations were initialized with equally abundant predators and prey, i.e. *C*_*i*_ = *R* = 0.1. We determined species coexistence based on the presence or absence of long-term trends in the population dynamics. First, for each predator, we calculated the *Pearson’s correlation coefficient* (*ρ*) between the predators’ abundance and time using the last 5000 time units of the corresponding simulation runs (cf. Klauschies et al. 2016). We then considered the two predators to persist when both correlation coefficients were non-significant, i.e. *ρ*_*i*_ ≈ 0. In contrast, when one of the two predators exhibited a significant negative correlation coefficient, it is assumed to experience competitive exclusion.

Finally, we estimated the absolute (*O*_*a*_) and the relative (*O*_*r*_) range of realized species coexistence as follows:

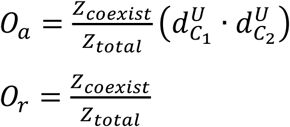

with *Z*_*coexist*_ being the number of simulations where the two predators coexisted. Simulations and analyses were performed in MATLAB, version 7.13, using solver ode23 for ODEs (The MathWorks Inc., Natick, MA, 2011). We increased the precision of the solver by reducing the absolute and relative tolerances to 10-9 and 10-11, respectively, and the maximum step size to 0.1.

## 3. Results

We investigated the coexistence of two predators on one prey through differences in the curvature of their functional responses and endogenously driven fluctuations in the prey abundance. The outcome mainly depends on the death rates of the two predators 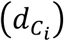 when their functional responses are locked (Abrams and Holt 2002). Hence, we evaluated the region in the two-dimensional 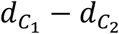 parameter space that may allow predator coexistence (Figure 2A; grey shaded area). The absolute size of this parameter region (*P*_*a*_) measures the overall strength of the corresponding coexistence mechanism. In contrast, the relative size of this parameter region 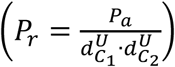 scales *P*_*a*_ with respect to 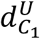 and 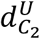, i.e. the upper limits of the predators’ death rates, and quantifies the proportion of the parameter space of predator persistence that may allow predator coexistence. Throughout the study, we assumed that predator one (*C*_1_) exhibits a linear functional response and predator two (*C*_2_) a type II functional response (Figure 2B).

Depending on the predator’s death rates, either both predators coexist on one prey due to endogenously generated predator-prey oscillations (Figure 2C) or one predator is competitive excluded by the other predator (Figure 2D-F). When *C*_1_ is the only remaining predator, the predator-prey dynamics approach a stable equilibrium over time because of its linear functional response (Figure 2D). In contrast, when the predators’ death rates select for *C*_2_, the corresponding predator-prey dynamics either show stasis (Figure 2E) or a limit cycle (Figure 2F).

To clearly relate our results to previous studies we first considered the classical predator-prey model where two predators are feeding on one prey that exhibits logistic growth in the absence of predation (Hsu et al. 1978; Abrams and Holt 2002; Xiao and Fussmann 2013). Hence, we first evaluated how *P*_*a*_ and *P*_*r*_ depend on the feeding traits, i.e. attack rates and handling times, of the different predators. For this, we set the corresponding parameters *h*_*R*_, 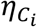, and *δ* of eq. [9] all to zero. After having established this baseline for further analysis, we investigated the impact of nutrient retention by the predators 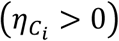, and of a non-linear nutrient uptake rate of the prey (*h*_*R*_ > 0) on *P*_*a*_ and *P*_*r*_ in the presence or absence of a shared loss rate (*δ* ≥ 0) simulating e.g. batch or chemostat conditions.

### 3.1 Relative non-linearity in the predators’ functional responses fosters coexistence

Predator coexistence on one prey essentially depends on the curvature of the type-II functional response of *C*_2_. It has to be sufficiently non-linear for the ecologically feasible prey densities in order to allow predator-prey oscillations. Therefore, a necessary requirement for predator coexistence is that the attack rate 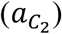 and the handling time 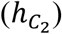 of *C*_2_ are sufficiently high, i.e. 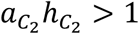 (for details see Appendix A). In addition, stable predator coexistence strongly relies on differences in their functional responses and thus, on deviations in their attack rates and handling times.

The absolute size (*P*_*a*_) and the relative size (*P*_*r*_) of the parameter region that may allow predator coexistence increase with 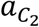 while keeping the attack rate of 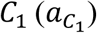 constant (Figure 3A-B). This happens because higher values of 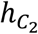 strongly increase the competitive ability of *C*_2_ compared to *C*_1_ at lower prey densities *R*, while hardly affecting its competitive ability at higher densities of *R*, thereby preventing the competitive exclusion of *C*_2_ by *C*_1_. Regarding the handling time 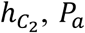 shows a humped-shape pattern, independent of the exact value of 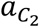 (Figure 4A): when starting from low values, an increase in 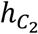 leads to an increase in *P*_*a*_ up to a value of 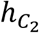 where *P*_*a*_ reaches a maximum. Increasing 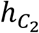 further causes *P*_*a*_ to decrease again due to the ambivalent effects of higher values of 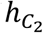 on the predator-prey dynamics. An increase in 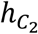 generally promotes predator-prey oscillations and thereby predator coexistence but it also reduces the maximum grazing rate of *C*_2_ and thus, the maximum death rate it can withstand. Consequently, depending on the net-effect of both processes, an increase of 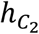 can lead either to an increase or a decrease of *P*_*a*_. This strongly contrasts with the always beneficial impact of 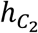 on *P*_*r*_ that is scaled with respect to the maxima of the predators’ feasible death rates (Figure 4B).

**Figure 3:**
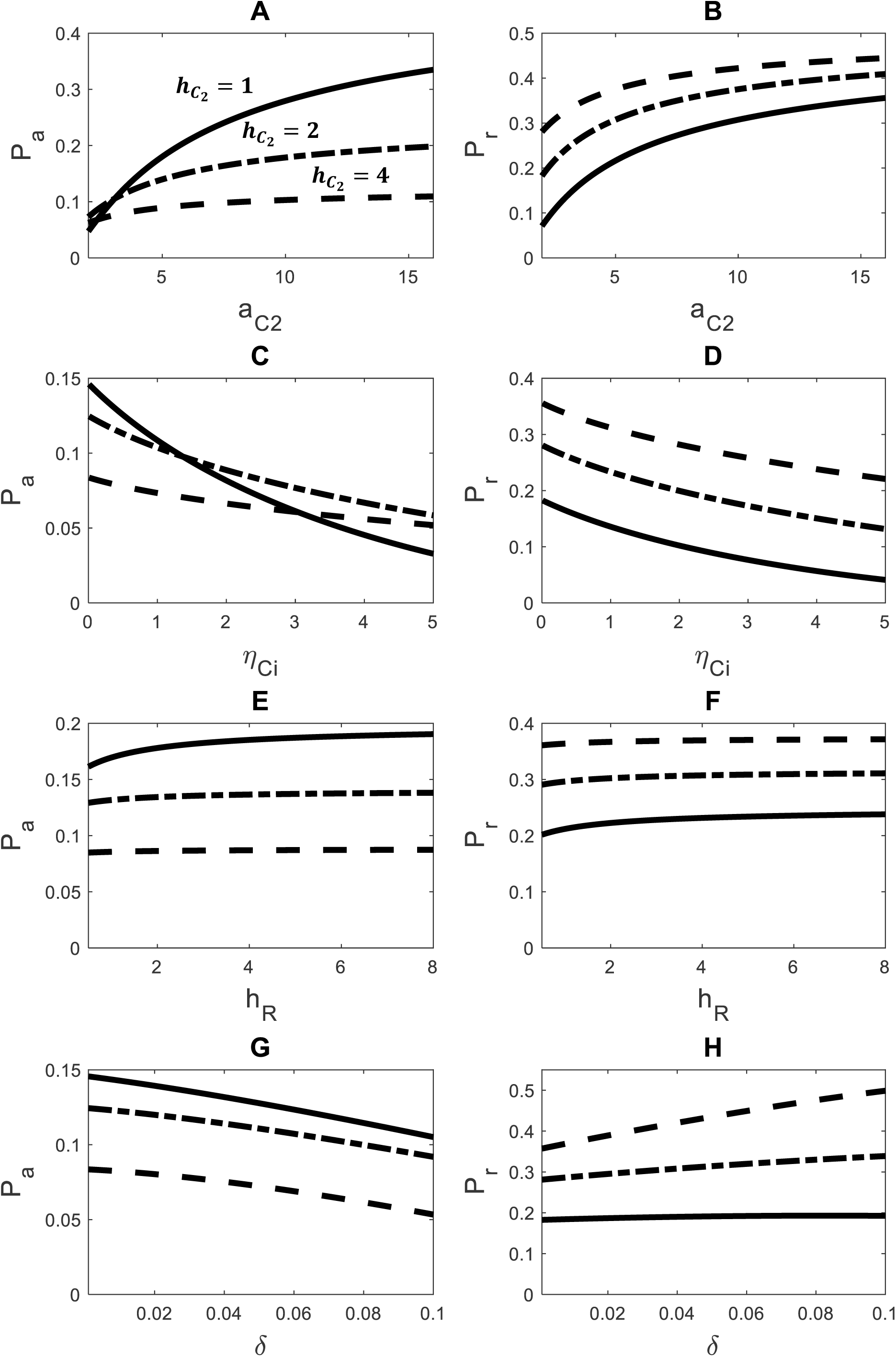
The absolute size, P_a_, (A, C, E, G) and the relative size, P_r_, (B, D, F, H) of the parameter range that may allow predator coexistence in dependence of the attack rate of *C*_2_, 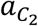 (A, B), the strength of nutrient competition between the predators and the prey, 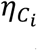, (C, D), the non-linearity of the prey’s nutrient uptake rate, h_R_, (E, F) or the shared loss rate for all species, δ (G, F) for three different values of the handling time of *C*_2_ (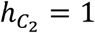, solid line;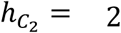, dashed-dotted line;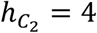, dashed line). Higher values of P_a_ indicate a larger region in the parameter space that may enable predator coexistence. The attack rate of *C*_1_, 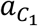, was set to 1. Increasing 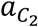 enhances the relative non-linearity between the functional responses of the two different predators. The higher 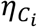, the more nutrients are stored in the biomasses of the predators and, thus, the higher is the competitive effect of the predators on the prey. Increasing δ enhances the overall mortality of all species which is most relevant for the persistence of *C*_2_. All results are based on the analytically derived necessary conditions of species coexistence (for details see Methods and Appendix A).

**Figure 4:**
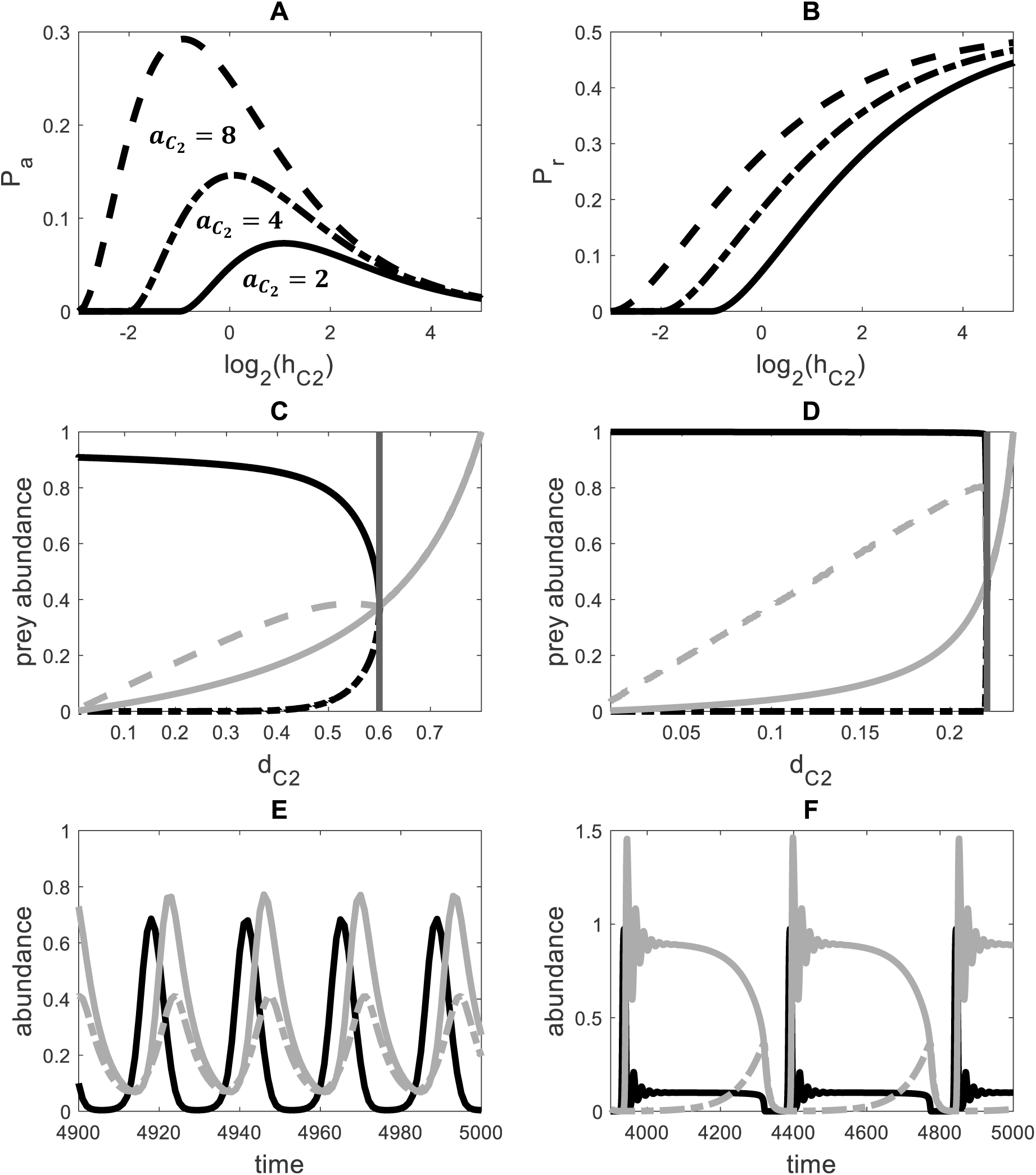
Panels A and B show the absolute size, P_a_, and the relative size, P_r_, of the parameter region that may allow predator coexistence in dependence of the handling time of *C*_2_, 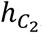, for three different values of the attack rate of *C*_2_ (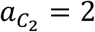, solid line;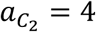, dashed-dotted line; 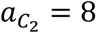, dashed line). Higher values of P_a_ indicate a larger region in the parameter space that may enable predator coexistence. Panels C and D show the maximum (solid black line), minimum (dashed-dotted black line), temporally averaged (dashed grey line) and equilibrium value (solid grey line) of the prey’s abundance (R) in dependence of the death rate of 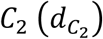 for the R − *C*_2_ system 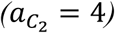, in case of 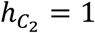(C) and 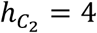 (D). The solid vertical grey line indicates the Hopf-bifurcation point 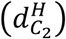 of the corresponding R − *C*_2_ system. Panels E and F display the population dynamics of R (black solid line), *C*_1_ (grey solid line) and *C*_2_ (grey dashed-dotted line). The shape of the predator-prey oscillations depends on the attack rate and handling time of *C*_2_ and the death rates of the two different predators, ranging from simple predator-prey oscillations 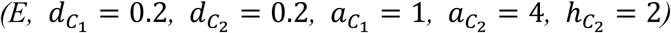 to complex population dynamics 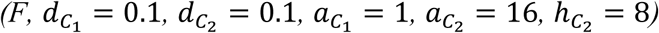.

Comparing *P*_*a*_ with the size of the parameter region where the predators definitely coexist (*O*_*a*_), for different values of 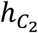, further reveals that *P*_*a*_ strongly overestimates *O*_*a*_ when 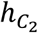 is small, but is of similar magnitude as *O*_*a*_ when 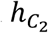 is sufficiently high. For example, for 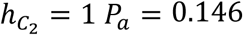 is more than twice as large as *O*_*a*_ = 0.062 (Figure 5A) but for 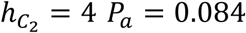 is nearly as large as *O*_*a*_ = 0.073 (Figure 5B). This happens because the amplitude of the predator-prey oscillations of the *R* − *C*_2_ system increases much slower as 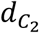 decreases from its Hopf-bifurcation point 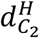 when 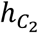 is small (Figure 4C, D). As a result, the difference between the temporal average of 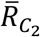 and its equilibrium value 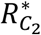 is substantially lower for 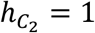 than for 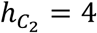 (Figure 4C, D). Hence, for 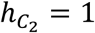, i.e. when the non-linearity of the type II functional response of *C*_2_ is rather weak, *C*_1_ cannot take advantage of very long periods of high resource densities, reducing its potential to invade the *R* − *C*_2_ system. Another consequence of this property is that the value of 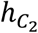 that maximizes *O*_*a*_ will be generally higher than the value that maximizes *P*_*a*_.

**Figure 5:**
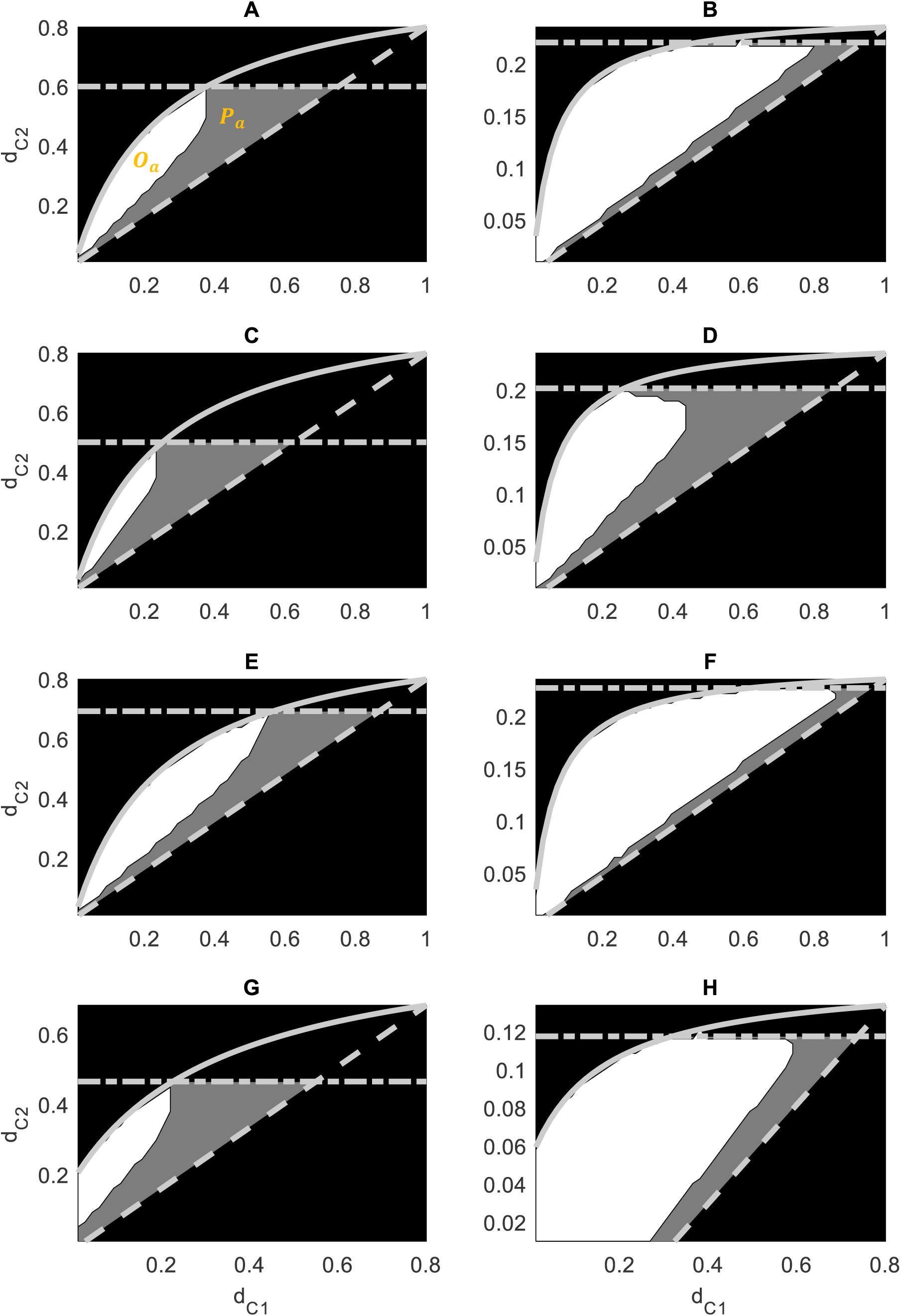
The outcome of predator competition in dependence of the death rates 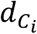 of two different predators *C*_1_ and *C*_2_ assuming logistic growth for the prey (A, B; h_R_ = 0, 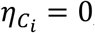, δ = 0), nutrient competition between predators and prey (C, D; h_R_ = 0, 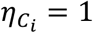, δ = 0), a non-linear nutrient uptake of the prey (E, F; h_R_ = 2, 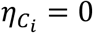, δ = 0) or an overall loss rate for all species (G, H; h_R_ = 0, 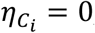, δ = 0.1). Results are shown for a lower (A, C, E, G;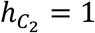) and a higher (B, D, F, H;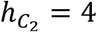) handling time of *C*_2_. The attack rates of the two different predators were set to 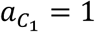 and 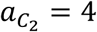. The parameter region that may potentially enable predator coexistence, P_a_, is indicated by the grey shaded area whereas the parameter region for which the two predators do actually coexist, O_a_, is marked by the white area. The black area indicates parameter combinations for which one of the two predators goes extinct (for further details see the caption of Figure 1).

Finally, when the two predators are able to coexist, the shape of the predator-prey oscillations can differ greatly, depending on the parameters of the functional responses and the death rates of the two predators (Figure 4E-F). For most parameter combinations, the oscillations are quite regular, typically exhibiting one dominant time scale at which the two predators are almost completely synchronized, i.e. their phase difference is close to zero (for an example see Figure 4E; further details are given in Appendix C). However, the amplitudes of the predators differ. *C*_1_ strongly increases when the prey density is sufficiently high, but also strongly decreases when the prey density is falling below its minimum prey requirement, promoting a rather high temporal variation in its abundance (*Coefficient of variation* (CV) = 0.77). In contrast, *C*_2_ increases less in response to high prey densities due to its lower maximum growth rate but also decreases less when the prey density is rather low because of its lower death rate and thus, higher starvation resistance at food shortage. As a result, the abundance of *C*_2_ often varies less (CV = 0.60). This type of dynamics nicely reflects the gleaner-opportunist trade-off faced by the two predators, where *C*_1_ exhibits a competitive advantage at times of high prey densities whereas *C*_2_ is doing better at periods of low prey densities.

This pattern strongly contrasts with the dynamics observed for some other parameter combinations, where the shape of the predator-prey oscillations is much more complex, often exhibiting two dominant time scales (Figure 4F). This type of dynamics is only observed when the non-linearity of the type II functional response of *C*_2_ is very strong. As a result of temporal niche differentiation between the two predators, the abundances of *C*_1_ and *C*_2_ vary asynchronously on the longer time scale, displaying a phase difference of 0.74 π (for details see Appendix C). In contrast, the predator-prey oscillations on the shorter time scale are only clearly visible in the dynamics of *R* and *C*_1_, preventing any meaningful quantification of the phase difference between the predators’ population dynamics. In line with the core mechanism that allows species coexistence through relative non-linearity in the predators’ functional responses, a temporary dominance of *C*_1_ strongly stabilizes the predator-prey oscillations. Under these conditions, *C*_2_ exhibits positive net-growth rates due to its lower prey requirements, allowing its regrowth from low densities. The enhanced grazing pressure by *C*_2_ then causes a decrease in *R* below the minimum prey requirement of *C*_1_ leading to a strong decrease in its abundance. The associated release from competition, in turn, allows *C*_2_ to establish high abundances. As a consequence of the curved functional response of *C*_2_ and its temporary dominance, the predator-prey dynamics start to exhibit fast predator-prey oscillations. As a result, the corresponding high mean values in *R* (immediately) favor the regrowth of *C*_1_ compared to *C*_2_. This induces low abundances of *C*_2_ again, resetting the initial conditions of the competitive cycle.

### 3.2 Nutrient retention by the predators hampers coexistence

Predator coexistence on one prey also depends on the type of resource limiting prey growth. Compared to light or space limitation (Figures 3A-B, 5A-B), nutrient limitation of the prey strongly reduces the parameter region for which the two predators coexist (Figures 3C-D, 5C-D) because it stabilizes the predator-prey dynamics (Appendix A). The ecological mechanism behind this stabilizing effect is that predators absorb mineral nutrients in their biomasses that are not further available for prey uptake. Hence, high predator densities do not only negatively affect the prey directly by predation, but also indirectly through nutrient retention. Therefore, the carrying capacity of the prey is effectively reduced in the presence of the predators, which is known to stabilize predator-prey dynamics.

The negative impact of nutrient retention on the absolute (*P*_*a*_) and relative (*P*_*r*_) sizes of the parameter range that may enable predator coexistence is more severe for lower values of 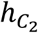 (Figures 5C-D vs. A-B). For example, *P*_*a*_ decreases by roughly 26% for 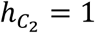 (Figure 5A&C) whereas the decrease is only about 13% for 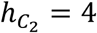 (Figure 5B&D). The negative impact of nutrient retention on predator coexistence is even more pronounced when considering the parameter range of realized predator coexistence (*O*_*a*_), especially for higher values of 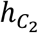. In fact, nutrient retention reduces the size of the parameter range of *O*_*a*_ by nearly 55% for 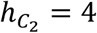 (Figure 5A&C) and by 44% for 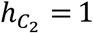 (Figure 5B&D). Hence, the impact of nutrient retention on predator coexistence is very pronounced for a broad range of parameters.

### 3.3 Non-linear resource uptake rates of the prey promote predator coexistence

Increasing the non-linearity of the resource uptake rate of the prey slightly benefits predator coexistence based on relative non-linearity in their functional responses (Figures 3E, F; 5E, F) because it promotes unstable predator-prey dynamics, especially for lower values of 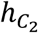. This can be seen by the higher values of the Hopf-bifurcation point 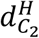 (dashed dotted lines in Figure 5E, F) at which the *R* − *C*_2_ system gives rise to predator-prey oscillations. In our example, a non-linear nutrient uptake rate increases the size of *P*_*a*_ by 18% and of *O*_*a*_ by 48% for 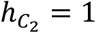 (Figure 5A&E). In contrast, the positive effect of a non-linear nutrient uptake rate is less pronounced for 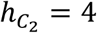, in which case the increase was limited to 2% for *P*_*a*_ and 8% for *O*_*a*_ (Figure 5B&F). This observation can be explained by the fact that *P*_*a*_ and *O*_*a*_ were already close to their maxima for 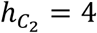, i.e. the Hopf-bifurcation point 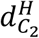 of the *R* − *C*_2_ system almost coincides with the upper limit of 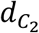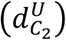 (cf. Figure 5B&F). In contrast, for 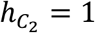, the positive effect of a non-linear nutrient uptake rate of the prey is quite large given that 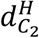 was substantially below 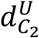 (cf. Figure 5A&E).

### 3.4 Coexistence is more likely in closed than in flow-through systems

Adding a shared loss rate to all species by considering a chemostat (i.e. flow-through) system with a dilution rate *δ* substantially reduces the possibility of predator coexistence on one prey. An increase in *δ* always strongly decreases the size of *P*_*a*_ and *O*_*a*_, independent of 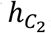, i.e. the exact curvature of the type-II functional response of *C*_2_ (Figures 3G, 5G-H vs. 5A-B). For example, *P*_*a*_ and *O*_*a*_ are reduced by 28% and 42%, respectively, when 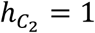 and by 36% and 40% when *h*_2_ = 4. This happens because *δ* reduces the preys’ carrying capacity, which stabilizes the predator-prey dynamics (Appendix A), and decreases the maximum death rates of the two predators they can withstand (cf. scale of y axes in Figure 5).

In contrast to *P*_*a*_, the relative size of the parameter region that potentially allows predator coexistence (*P*_*r*_) increases with *δ*, independent of 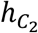 (Figure 3H). For instance, *P*_*r*_ increases by 6% when 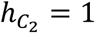 and by 29% when *h*_2_ = 4 (Figure 5G-H vs. 5A-B). This is explicable by the larger negative impact of *δ* on *C*2 than *C*1 due to its lower maximum growth and death rates, which renders *C*_2_ more sensitive to additional mortality (cf. Figure 2A, B). Consequently, *C*_2_ cannot exclude *C*_1_ anymore when the death rate of *C*_1_ is sufficiently low, e.g. below ≈ 0.3 in Figure 5H (dashed line). Furthermore, *P*_*r*_ may exceed 0.5 when the dilution rate is sufficiently high (Figure 3H), which strongly contrasts with closed systems where *δ* = 0.

In contrast to *P*_*a*_, *O*_*a*_ and *P*_*r*_, the response of the relative size of the parameter region that actually allows predator coexistence (*O*_*r*_) to alterations of *δ* depends on 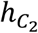. For example, *O*_*r*_ decreases by 14% when 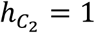 (Figure 5G vs. 5A) because the stabilizing effect of *δ* on the predator-prey dynamics reduces predator coexistence more than any beneficial effect arising through a reduced risk of *C*_1_ to get excluded by *C*_2_ at lower death rates of both predators. In comparison, *O*_*r*_ increases by 25% when *h*_2_ = 4 (Figure 5H vs. 5B) because the positive effect of *δ* on predator coexistence at lower death rates outweighs the weakly stabilizing effect of *δ* on the predator-prey dynamics and thus negative impact on predator coexistence.

## 4. Discussion

For almost 50 years, it has fascinated ecologists that two different predators may be able to coexist on one fluctuating prey when they exhibit substantial differences in the curvature of their functional responses (Koch 1974; Chesson 2000; Abrams and Holt 2002). Previous studies explored this mechanism of species coexistence in predator-prey models that are based either on the classical Rosenzweig-MacArthur (RM) equations (1963) with logistically growing prey (Hsu et al. 1978; Armstrong and McGehee 1980; Abrams 2004; Xiao and Fussmann 2013), or on chemostat equations with explicit nutrient dynamics (Butler and Waltman 1981; Butler et al. 1983; Keener 1985; Smith 1995). The main motivation for using the latter was to derive predictions, which can be experimentally tested (e.g. Butler and Waltman 1981). The RM equations are often believed to mainly differ from corresponding chemostat models by assuming a linear uptake rate of the prey for the limiting resource (e.g. Grover and Holt 1998). This impression arises because logistic growth, as it is modelled by RM equations, can be derived from an explicit consumer-resource model that assumes a linear uptake rate of the consumer for the limiting resource (Armstrong and McGehee 1980). However, this derivation is based on the important assumption that no other trophic levels, such as predators, are present. By adding another trophic level to the original system, we show that predators may influence their prey not only by predation but also by nutrient retention (Figure 1; Appendix A). Hence, the RM model reflects conditions where the prey is limited by a resource not affected by the predators such as light or space, rather than nutrients. The two different types of predator-prey models thus substantially differ in their assumption about the type, uptake and replenishment of the resource that is limiting the growth of the prey.

Therefore, we analyzed a generic predator-prey model that accounts explicitly for the uptake and replenishment of nutrients and compared it to the classical RM predator-prey model. We evaluated the impact of nutrient retention by the predators on their coexistence on one prey through relative non-linearity in their functional responses and associated fluctuations in prey abundance. Our results demonstrate that nutrient retention strongly stabilizes the dynamics and thereby substantially reduces the potential for predator coexistence. The stabilizing effect of nutrient retention originates from the negative impact of the predators on the effective carrying capacity of their own prey. Our study thus demonstrates that the type of resource limitation may substantially influence the stability of predator-prey dynamics and the likelihood for predator coexistence through fluctuation-dependent mechanisms, with potentially far-reaching consequences for the overall stability of entire food-webs.

Since nutrient limitation of the (autotrophic) prey is common in aquatic (Sommer and Lampert 2007) and terrestrial ecosystems (Santiago and Goldstein 2016), we can expect that predators very likely influence the amount of the limiting resource for their prey in many natural systems. This holds in particular for (aquatic) ecosystems that frequently exhibit top-heavy biomass distributions in which the heterotrophic biomass is at least of similar magnitude as the autotrophic biomass (del Giorgio and Gasol 1995; Burkepile 2013; McCauley et al. 2018; Shurin et al. 2006). Given the high nutrient to carbon ratios of animals compared to plants (Sterner and Elser 2002; Hall 2009; Hessen et al. 2013), the majority of nutrients in aquatic ecosystems is stored in the consumer biomass (Hassett et al. 1997; Gaedke et al. 2002). Hence, our predator-prey model described by eq. [9] captures an essential property of many natural systems, which is not entailed in the Rosenzweig-MacArthur (RM) equations (1963). This is particularly relevant, because important ecological features such as the paradox of enrichment (Rosenzweig 1971), the hydra effect (Abrams 2009; Sieber and Hilker 2012), biological chaos in three species food chains (Hastings and Powell 1991; McCann and Yodiz 1994), the impact of trait adaptation on species coexistence (Abrams 2006; Klauschies et al. 2016) and the stability of complex food webs (Brose et al. 2006; Williams and Martinez 2008) were investigated in models based on the RM equations.

Nutrient retention by predators may even be relevant for (terrestrial) ecosystems with bottom-heavy biomass distributions (Polis 1999; Shurin et al. 2006; Hatton et al. 2015). On a global scale, the biomass ratio of heterotrophs to autotrophs is about 0.05 in terrestrial ecosystems (Bar-On et al. 2018). Assuming a five times higher nutrient to carbon ratio for heterotrophs than for autotrophs (e.g. Lemoine et al. 2014; Sterner and Elser 2002), the heterotrophic biomass contains 20% of the total amount of nutrients. Hence, nutrient retention by heterotrophs is likely to be important also for the overall population dynamics in terrestrial ecosystems. Only if predators mainly affect their prey by consumption and less by nutrient retention, predator-prey models based on the Rosenzweig-MacArthur equations (1963) may well capture the observed predator-prey dynamics, e.g. in hyper-eutrophic systems where prey is rather light-than nutrient-limited (Hautier et al. 2009).

In line with previous work (DeAngelis 1992; Leroux and Schmitz 2015), our results demonstrate that it is very important to account explicitly for the nutrient dynamics in food web models because predators play an import role in the recycling and translocation of mineral nutrients in natural systems (Vanni 2002; Schmitz et al. 2010). Whereas nutrient retention acts as a stabilizing factor in predator-prey models (Appendix A), the process of nutrient recycling is destabilizing (Appendix D), which is in accordance with previous studies (Kooi et al. 2002; Kuijper et al. 2004; Quévreux et al. 2018). Hence, resolving the energetic and elemental pathways in ecological models may inform us about essential features of natural systems that may either promote or hinder the stability of entire food webs (cf. DeAngelis 1992; Kooijman 2000; Sterner and Elser 2002).

Our study further shows that predator coexistence on one prey also depends on the non-linearity of the prey’s uptake rate for the resource. A linear uptake rate is to be expected under strong bottom-up control of the prey where the limiting resource is scarce, e.g. severe light limitation in winter or ultra-oligotrophic conditions in the open ocean (Lampert and Sommer 2007; Sommer 2005). Otherwise non-linear relationships are more realistic (Michaelis and Menten 1913; Murdoch 1973). This is very important given that our results show that a non-linear nutrient-uptake rate may strongly promote predator-prey oscillations, and thereby facilitate predator coexistence when the relative non-linearity between the predators’ functional responses is relatively weak. In contrast, a non-linear resource uptake rate of the prey hardly influences the likelihood for predator coexistence when the relative non-linearity between the predators’ functional responses is rather strong. This is in line with previous findings from a modified RM predator-prey model that includes theta-logistic growth to reflect a non-linear nutrient uptake rate of the prey (Abrams and Holt 2002). Overall, predator coexistence is more or less common in our model compared to the corresponding RM model with logistically growing prey, depending on whether the stabilizing effect of nutrient retention dominates over the destabilizing effect of a non-linear nutrient uptake rate.

Finally, our results demonstrate that predator coexistence on one prey becomes more difficult if all species experience a shared loss rate because it reduces the effective carrying capacity of the prey, and the maximum death rates the predators can withstand. Furthermore, in line with previous results for standard chemostat models (Smith 1995), a higher dilution rate stabilizes predator-prey dynamics and thereby also reduces the likelihood for predator coexistence through relative non-linearity in their functional responses. While adding a shared loss rate to our predator-prey system does not alter the total amount of nutrients in the system, it does affect the distribution of nutrients, i.e. the ratio between nutrients bound in biomass and dissolved inorganic nutrients. A shared loss rate may naturally arise through water exchange or an omnivorous top predator feeding on predators and prey.

There are two fundamental requirements for predator coexistence through relative non-linearity. One are fluctuations in the abundance of the prey that can be externally driven (Sommer 1985; Grover 1988; Abrams 2004) or endogenously generated by predator-prey interactions (Armstrong and McGehee 1980). The latter essentially depends on the potential of the predators’ functional response to destabilize the dynamics, for which at least one functional response has to be sufficiently non-linear for the ecologically feasible prey densities. This condition is likely to be met in many natural systems, given ample experimental evidence for invertebrate predators to exhibit substantial curvature in their functional responses at relevant prey densities (Hassell et al. 1977; Rothhaupt 1988; Jeschke et al. 2004; Sarnelle and Wilson 2008; Seifert et al. 2014).

The second requirement for predator coexistence are differences in the predators’ abilities to exploit efficiently periods of low or high prey abundances in line with the gleaner-opportunist trade-off (Fredrickson and Stephanopoulos 1981; Xiao and Fussmann 2013). Experimental results and field observations suggest that this trade-off may result from size differences among predators or from differences in their feeding or life-history strategies. For example, smaller rotifers (the gleaners) had lower prey requirements than larger rotifers allowing the smaller species to outcompete the larger species at lower prey densities (Stemberger and Gilbert 1985; Sarma et al. 1996). In contrast, larger rotifers (the opportunists) are better competitors at higher prey densities, enabling them to exploit favorable conditions of high prey densities (Boraas et al. 1990; Sarma et al. 1999). Another example are filter-feeding zooplankton such as cladocerans with often linear functional responses (Jeschke et al. 2004) and high maximum growth rates (Sommer and Stibor 2002). In contrast, interception feeders such as copepods show non-linear functional responses (Jeschke et al. 2004) and lower maximum growth rates (Sommer and Stibor 2002).

However, a direct experimental demonstration of predator coexistence through endogenously generated fluctuations in prey abundance and differences in the functional responses of different predators is still lacking. An important reason might be that different coexistence mechanisms frequently operate in parallel and that it is rather difficult to experimentally test for the individual effect of a particular coexistence mechanism (Ellner et al. 2018). In fact, predator coexistence on one prey is also possible in the absence of predator-prey oscillations, for example, when predators show interference between their conspecifics (Vance 1984; Hsu et al. 2013), utilize different life stages of the prey (Haigh and Maynard Smith 1972), or are limited by different currencies of the prey such as food density and food quality (Loladze et al. 2000, 2004; Moe et al. 2005; Elser et al. 2012).

## Conclusions

We show that the coexistence of two predators on one prey through differences in the curvature of their functional responses and associated fluctuations in prey abundance depends on the type of resource limiting prey growth. In contrast to light or space limitation, predators retain part of the limiting resource in their biomass when the prey is subject to nutrient limitation, which reduces the availability to their prey. This feature strongly stabilizes the predator-prey interaction and thereby reduces the possibility for the two predators to coexist in a fluctuation-dependent way. In contrast, a non-linear resource uptake rate of the prey promotes predator-prey oscillations, and thus, predator coexistence. Furthermore, predator coexistence through relative non-linearity in their functional responses is less likely to occur in flow-through systems where all species suffer from an additional shared loss rate. Hence, fluctuation-dependent mechanisms of species coexistence are most likely to promote biodiversity in closed systems where the basal prey experiences light or space limitation. In contrast, under nutrient limitation, the maintenance of biodiversity may strongly rely on fluctuation-independent mechanisms.

## Author contributions

Toni Klauschies conceived the idea of this study, developed the mathematics included, coded and simulated the model, analyzed the data and wrote the manuscript. Ursula Gaedke commented on the manuscript and improved the writing.

## Acknowledgments

We thank Michael Sieber, Ruben Ceulemans, Alice Boit and two anonymous reviewers for helpful comments and suggestions. Toni Klauschies was funded by the German Research Foundation (DFG, GA 401/26-1).

## Appendix A Necessary conditions and parameter range of potential species coexistence

Following the general methodology of Hsu et al. (1978) and Xiao and Fussmann (2013) we here derive ecological conditions for species coexistence that determine a range of parameters for which species coexistence might be possible. Although these conditions are not sufficient, they are necessary, implying that outside this parameter range species coexistence is impossible. For simplicity, we will assume predator one (*C*_1_) to exhibit a linear functional response and predator two (*C*_2_) to have a (non-linear) functional type II response.

### 1) Persistence of predator-prey systems in the absence of exploitative competition

Predator coexistence necessarily requires persistence of the two different predator-prey systems comprising either *C*_1_ or *C*_2_ next to the prey. Hence, each predator has to be able to invade the system that only includes the prey. In the absence of *C*_1_ and *C*_2_, the prey is approaching its single-species equilibrium 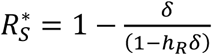. The two predators thus exhibit positive invasion growth rates when their grazing rates for 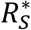 are larger than the sum of their different loss rates through mortality 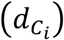 and dilution (*δ*), respectively, i.e. 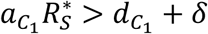 for *C*_1_ and 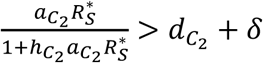 for *C*_2_. Consequently, the death rates that allow persistence of the two different predator-prey systems are limited to 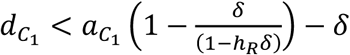 for *C*_1_ and to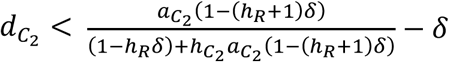 for *C*_2_. These conditions put natural limits on the two-dimensional parameter space in the 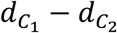 plane for which predator coexistence might be possible. The first necessary condition for predator coexistence is thus given by:

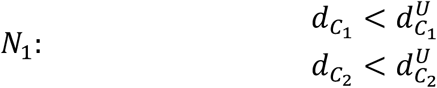

with 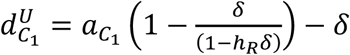 and 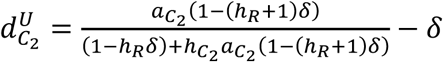 denoting the upper limits of the death rates of two different predators *C*_1_ and *C*_2_, respectively.

### 2) Invasion of *C*_2_

The linear functional response of *C*_1_ ensures that the *C*_1_ − *R* system will approach a stable equilibrium with prey density 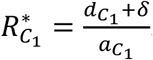. A necessary condition for predator coexistence is that *C*_2_ is able to invade the *C*_1_ − *R* system. Hence, the invasion growth rate of *C*_2_ for 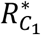 has to be larger than zero, i.e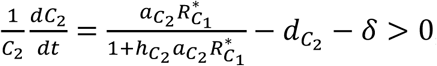, which corresponds to 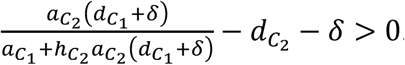. Therefore, the second necessary condition for predator coexistence is given by:

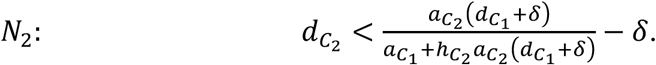

This implies that the equilibrium density of the prey in the *C*_2_ − *R* system has to be smaller than 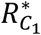, i.e.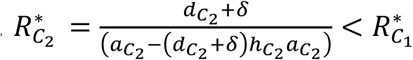. In other words, the minimum prey requirement of *C*_2_ has to be lower than the minimum prey requirement of *C*_1_. In contrast, when 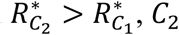 cannot invade the *C*_1_ − *R* system.

### 3) Competitive exclusion of *C*_1_

Since the curvature of the type II functional response is concave, the time-averaged per-capita net-growth rate of *C*_2_ will always be larger than or equal to the time-averaged per-capita net-growth rate of a corresponding predator that has a linear functional response with the same per-capita net-growth rates at the minimum (*R*_*min*_ = 0), and the maximum 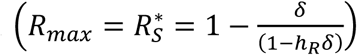 feasible prey densities as *C*_2_. Hence, the following inequality holds:

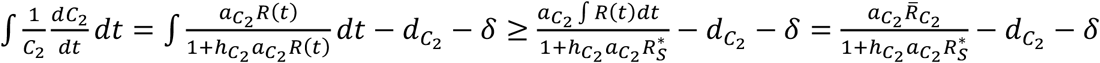

with 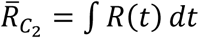 *dt* being the time-averaged prey density of the *C*_2_ − *R* system. Since 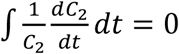 at equilibrium, the following inequality holds: 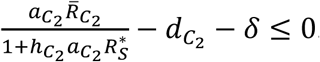 Setting the term 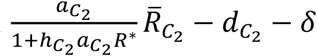 to zero and solving for 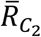, thus gives us an upper limit for the temporal average of the prey density in the corresponding predator-prey system that is denoted by 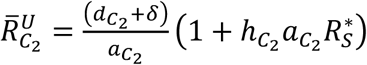. Therefore, whenever *C*_1_ has a minimum prey requirement 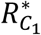 higher than 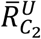, it will inevitably be outcompeted by *C*_2_. Consequently, the following inequality has to be met for the two predators to be able to coexist:

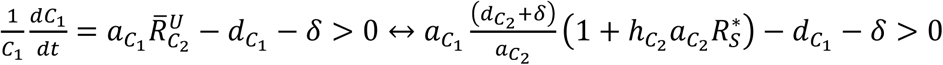

This leads to the following third necessary condition of predator coexistence:

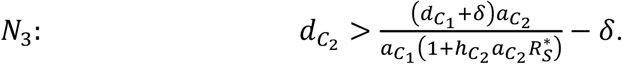

Nevertheless, 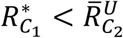 does not ensure that *C*_1_ can invade the *C*_2_ − *R* system. For this to happen, the minimum prey requirement of *C*_1_ has to be lower than the actual temporal average of the prey density in the corresponding predator-prey system, i.e.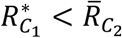.

### 4) Hopf-bifurcation in predator-prey models with type II functional response

Predator coexistence through relative non-linearity in their functional responses requires oscillations in the abundances of the different species. It is well known from the theory of non-linear dynamics that limit cycles only emerge in systems of ordinary differential equations with more than 2 dimensions when there is some degree of non-linearity, i.e. non-linear density-dependence, in the per-capita rates of the corresponding state variables. Hence, in the predator-prey system defined by eq. [9], oscillations can only occur in the presence of *C*_2_. Therefore, we performed a linear stability analysis of the two-dimensional *R* − *C*_2_ system, in order to evaluate the influence of nutrient retention by the predators, the effect of a non-linear resource uptake rate of the prey, and the impact of a shared loss rate for all species on the stability of the predator-prey dynamics. For simplicity, we will neglect the index of predator two for its state variable and parameters and thus consider the following predator-prey model:

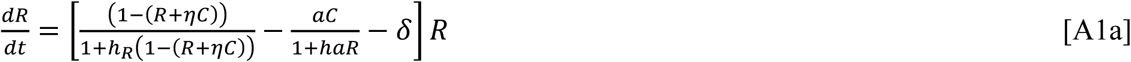

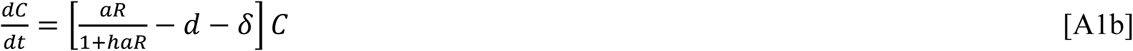

Throughout the analysis, we will focus on parameter values where the predator is able to persist, i.e. *d* < *d*^*U*^. To start, we calculate the interior equilibrium of the prey (*R*^*^) and predator (*C*^*^) species for the different model parametrizations used in the main text by setting eq. [A1] to zero, i.e. 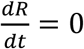 and 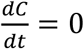, and solving for the corresponding values of the state variables satisfying these two conditions (for the calculated equilibria see Table A1). Subsequently, we will calculate the entries of the Jacobian matrix of the linearized predator-prey system and evaluate them at the interior equilibrium in order to describe the local behavior of the predator-prey system closely around its equilibrium state, and to determine its local stability. The stability of the interior equilibrium depends on the properties of the Jacobian matrix:

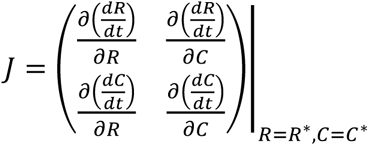

The predator-prey model defined by eq. [9] (main text) does not account for interference or cooperation among individual predators or any other kind of direct density-dependence in *C*. Consequently, the lower right entry of *J* is zero, i.e.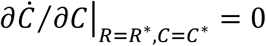. Furthermore, an increase in the density of the prey has always a positive effect on the predator density due to its type II functional response, which is a monotonically increasing function in *R*. Hence, the lower left entry of *J* is strictly positive, i.e. 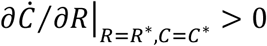. In contrast, an increase in the predator density will always have a negative effect on the prey density through grazing and nutrient retention. Therefore, the upper right entry of *J* is strictly negative, i.e. 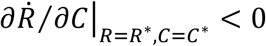. Finally, the sign of the upper left entry of *J*, i.e. 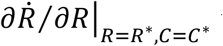 will depend on the actual parameter values, e.g. the death rate of the predator. Hence, the entries of *J* possess the following signs:

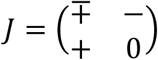

According to the Ruth-Hurwitz criterium for a two-dimensional system of ordinary differential equations, the interior equilibrium is stable when the determinant of the Jacobian matrix evaluated at the interior equilibrium (*J*) is positive and the corresponding trace is negative (Murray 2002). The determinant of the Jacobian matrix is given by:

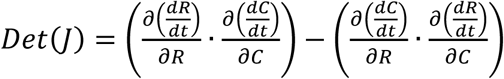

In line with our reasoning above, the sign of *Det*(*J*) is as follows:

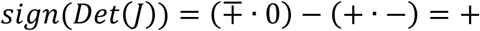

Therefore, the determinant of the Jacobian will always be positive. Consequently, the stability of the interior equilibrium entirely depends on the trace of the Jacobian matrix, which is given by:

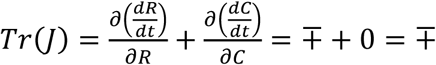

Since 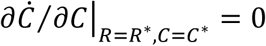, the trace of the Jacobian is given by the derivative of the rate of change of the prey with respect to its own density. Hence, we calculated 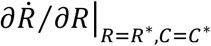 for the different model parametrizations used in the main text, and generally solved for the death rate at which the trace of the Jacobian is zero. At this particular point (*d*^*H*^), a Hopf-bifurcation occurs, separating two regions for which the system exhibits either a stable equilibrium (*d* > *d*^*H*^), or a limit cycle (*d* < *d*^*H*^) (cf. Figure 4C, D). In general, the Hopf-bifurcation point *d*^*H*^ has to be calculated numerically. However, in some cases explicit expressions for 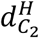 can be obtained (for details see Table A1).

**Table A1:**
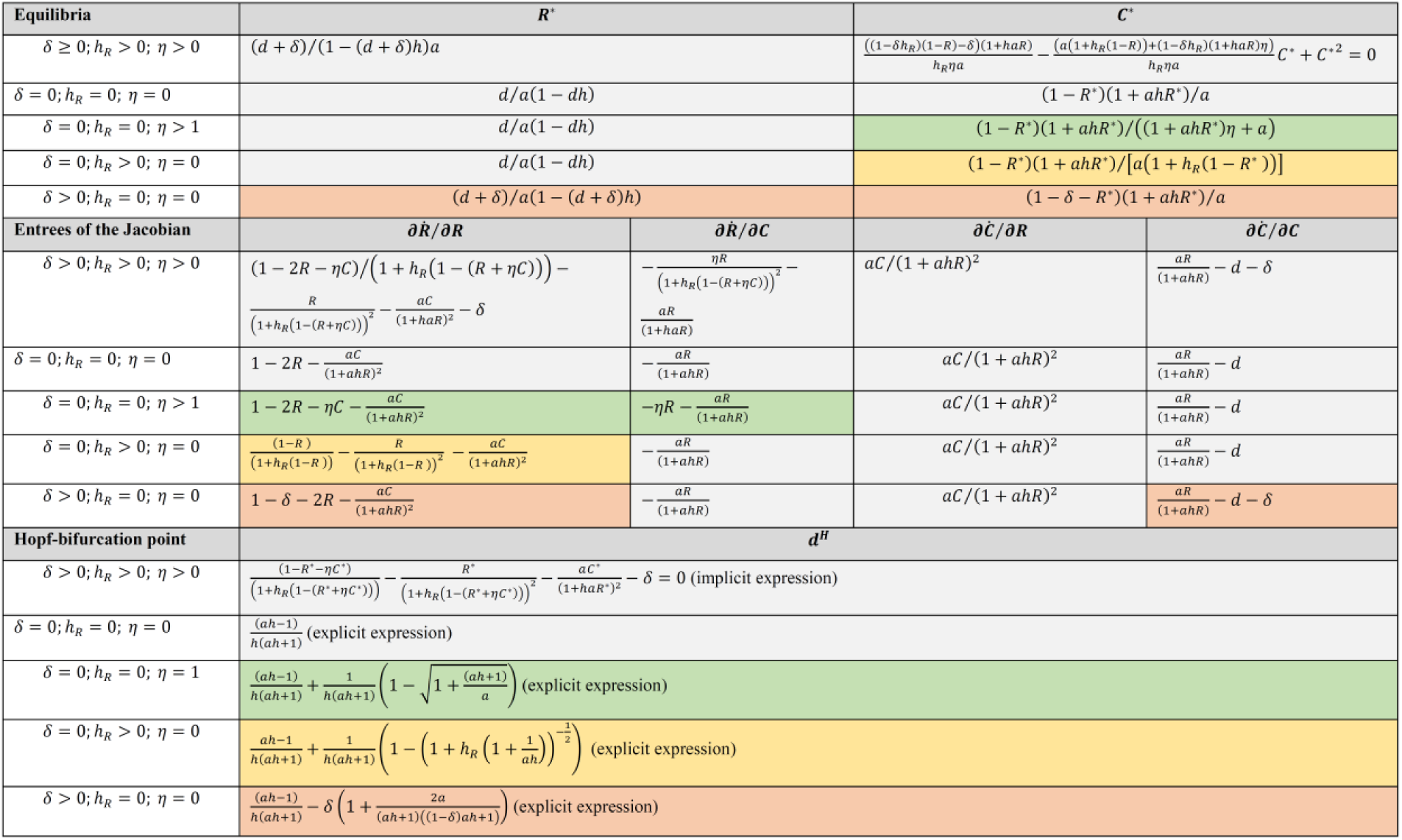
Summary of the linear stability analysis of the predator-prey model defined by eq. [A1] for the interior equilibrium under different model parameterizations, showing the influence of different ecological processes on the stability of the corresponding predator-prey dynamics. The colors highlight instances where the calculated terms of the particular expressions, i.e. equilibrium values, entrees of the Jacobian Matrix or Hopf-bifurcation point, differ from the ones of the reference scenario, i.e. the terms of the Rosenzweig-MacArthur predator-prey model (Rosenzweig-MacArthur 1963).

For instance, in the absence of nutrient retention by the predators (*η* = 0) and in case of a linear resource uptake rate of the prey (*h*_*R*_ = 0) and batch culture conditions (*δ* = 0) the Hopf-bifurcation point *d*^*H*^ of the classical predator-prey system (Rosenzweig and MacArthur 1963) with logistically growing prey in the vacancy of the predator is given by the following (see also Abrams and Holt 2002):

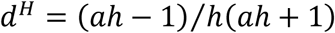

Since *d* > 0, a Hopf-bifurcation only occurs when *ah* > 1, so that *d*^*H*^ is also positive.

Furthermore, nutrient retention by the predators generally reduces the Hopf-bifurcation point of the corresponding *C* − *R* system and thus stabilizes its population dynamics. This can be seen when choosing a particular value for the competition coefficient *η* between the predator and the prey, e.g. *η* = 1. In this case, the Hopf-bifurcation point *d*^*H*^ can be written as:

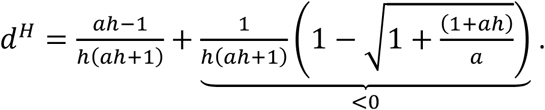

The stabilizing effect of nutrient retention on the predator-prey dynamics is stronger, when the non-linearity of the type II functional response is weaker. In contrast to the stabilizing effect of nutrient retention by the predators, a non-linear resource uptake rate of the prey has a destabilizing effect on the predator-prey dynamics. When accounting for non-linearity in the preys’ resource uptake rate (*h*_*R*_ > 0), the Hopf-bifurcation point of the corresponding predator-prey system is given as follows:

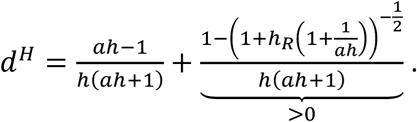

Hence, *d*^*H*^ increases with an increase in *h*_*R*_. The impact of *h*_*R*_ on the overall stability of the predator-prey dynamics depends on the curvature of the type-II functional response of the predator. The larger the product *ah* is, the larger the factor 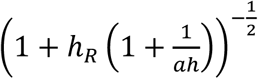 becomes and thus, the smaller the term 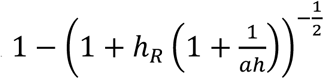 will be. Hence, the destabilizing effect of *h*_*R*_ on the predator-prey dynamics is larger when the non-linearity of the type-II functional response of the predator is weaker.

Finally, a common death rate of all species through the dilution rate *δ* is stabilizing the predator-prey dynamics. The Hopf-bifurcation point of the corresponding *C* − *R* system, i.e. when *δ* > 0, is given by the following:

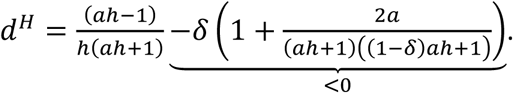

Hence, *d*^*H*^ decreases when *δ* is increasing. The stabilizing effect of *δ* on the predator-prey dynamics is larger when the non-linearity of the type-II functional response of the predator is stronger, and thus, when the handling time *h* is higher. This happens, because the dilution rate does not only affect the stability of the predator-prey dynamics but also the persistence of the corresponding *C* − *R* system.

### 5) Quantifying the size of the parameter range of potential species coexistence

The necessary conditions *N*_2_, *N*_3_ and *N*_4_ of predator coexistence derived above determine a range of parameters that may permit the coexistence of the two different predators (Figure A1A). In order to quantify the size of this parameter region we first calculated three different threshold values of the death rate of 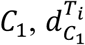, that allows piecewise integration of the relevant functions. The first threshold value 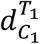 is given by the death rate 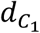 for which the function 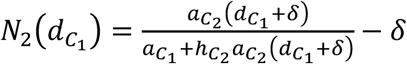 (reflecting the second necessary condition) intersects with the Hopf-bifurcation point 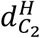 (denoting the fourth necessary condition), i.e. 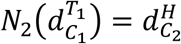 (cf. Figure A1B). Solving this equation with respect to 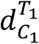 leads to the following expression of the first threshold value:

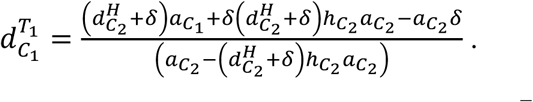

Accordingly, the second threshold value 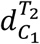 is given by the intersection point of the function 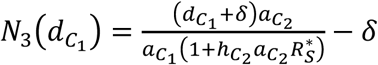 (representing the third necessary condition) with the Hopf-bifurcation point 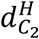, i.e. 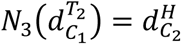 (cf. Figure A1C). Evaluating this equation with respect to 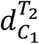 results into the following expression of the second threshold value:

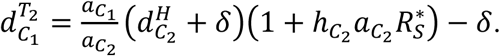

Finally, the third threshold value 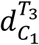 is given by the the value of 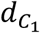 at which the function 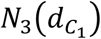 equals zero, i.e. 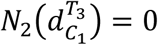 (cf. Figure A1D). Solving the latter for 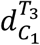 results into the following expression of the third threshold value:

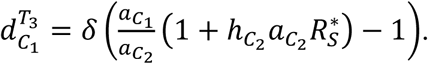

Using these three different threshold values, the absolute size of the parameter region of potential predator coexistence is given by the following sum of integrals (cf. Figure A1):

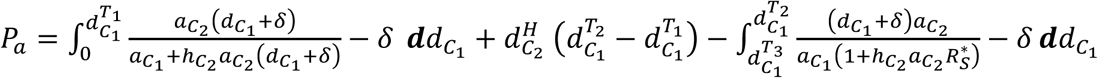

After having solved the first integral with *Wolfram Alpha*, we obtain the following expression of *P*_*a*_:

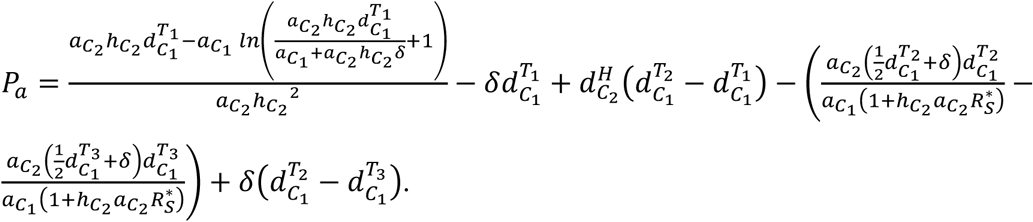

Note, although it is not explicitly stated, *P*_*a*_ also depends on 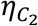 through the dependency of the Hopf-bifurcation point 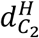 on 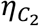. However, *P*_*a*_ does not depend on 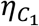. This strongly contrasts with the fact that the parameter range of realized species coexistence will likely also depend on 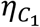 and potential differences between 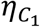 and 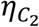.

**Figure A1:**
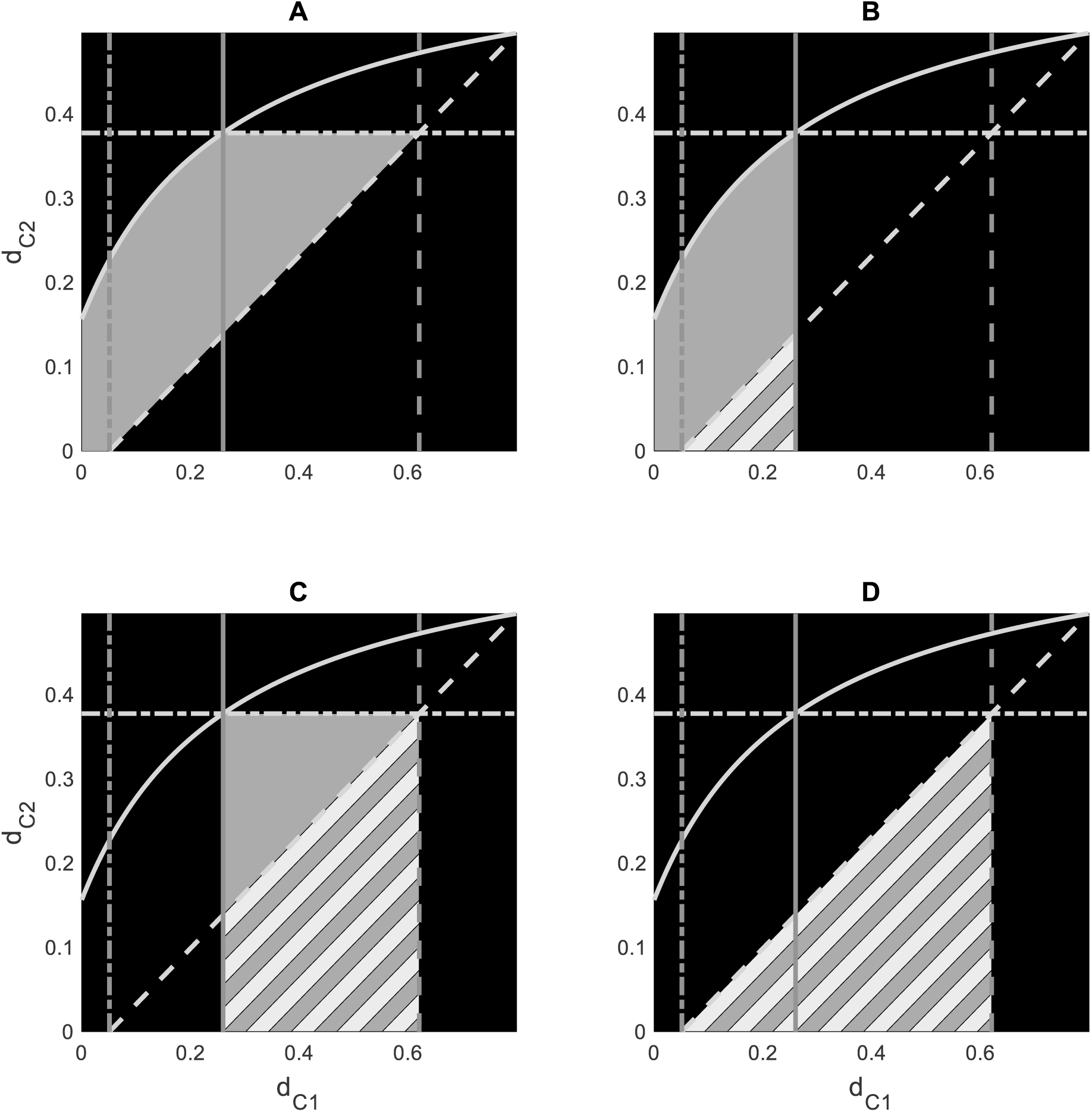
Parameter range, i.e. death rates 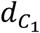 and 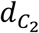, that potentially enables coexistence of predator one, *C*_1_, and predator two, *C*_2_ (grey shaded area) (for details see legend of Figure 1). The parameter range of potential species coexistence marked in panel (A) is given by the sum of parameter ranges shown in panels (B) and (C) minus the parameter range shown in panel (D). Striped areas in panels (C)-(D) indicate parameter regions where predator coexistence is impossible.

## Appendix B: Fluctuations in the prey abundance act as a stabilizing mechanism of predator coexistence

Contemporary theory suggests that species coexistence depends on the balance between stabilizing niche differences and destabilizing fitness differences among species (Chesson 2000; Barabás et al. 2018). Niche differences enable species coexistence by increasing intra-relative to interspecific competition, thereby facilitating the regrowth of species from low densities (Chesson 2018). In contrast, fitness differences promote competitive exclusion of inferior species in the absence of niche differentiation (Letten et al. 2017). Hence, two or more predators cannot coexist on one prey in the absence of niche differences, because the presence of inevitable fitness differences among the different predators will drive all but one of them extinct (Chesson 2000). Stabilizing niche differences between two different predators may arise through relative non-linearity in their functional responses and associated fluctuations in the prey abundance. We therefore here derive corresponding terms reflecting average fitness differences and stabilizing niche differences in our predator-prey model defined by eq. [9]. Following Chesson (2018), average fitness differences between the different predators can be inferred from their invasion growth rates in the absence of stabilizing niche differences. Hence, we calculated the invasion growth rates, i.e. temporal averages of the per-capita net-growth rates, of both predators for the corresponding resident system that comprises the other predator and the prey at its long-term equilibrium, which is assumed to be stable. Hence, the invasion growth rate of predator one for the equilibrium density of the prey in the corresponding resident predator-prey system comprising predator two is given by:

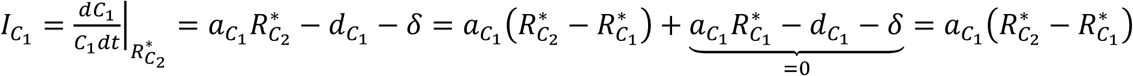

Similarly, the invasion growth rate of predator two for the equilibrium density of the prey in the corresponding resident predator-prey system comprising predator one is given by:

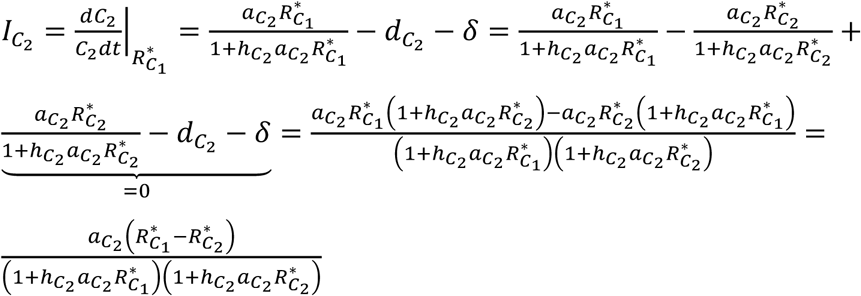

Comparing the invasion growth rates of both predators reveals that the competitive outcome entirely depends on the difference between 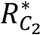 and 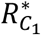. Hence, the average fitness differences (FD) between the two predators are given by the difference between the prey requirements of predator two 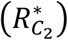 and predator one 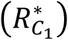, i.e.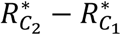.

According to the coexistence mechanism of relative non-linearity in the predator’s functional responses, stabilizing niche differences between the two different predators only arise through predator-prey oscillations. Given that the linear functional response of predator one ensures that the equilibrium of the corresponding predator-prey system is always stable, additional terms cannot arise in the invasion growth rate of predator two. However, the type-II functional response of predator two may destabilize the equilibrium of the corresponding predator-prey system. Under such conditions, the invasion growth rate of predator one is given as follows:

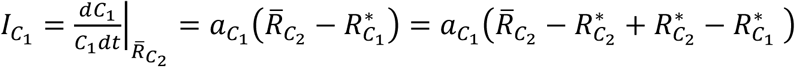

Comparing the invasion growth rate of predator one for the resident system of predator two in the absence of predator-prey oscillations, which prevents coexistence, with the one in the presence of predator-prey oscillations, which potentially allows coexistence, shows that the stabilizing niche differences (ND) between the two predators are given by the difference between 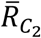 and 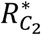, i.e.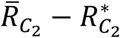. Hence, the net-stabilizing effect of relative non-linearity in the predators’ functional response on predator coexistence is thus given by the sum of stabilizing niche differences (+) and destabilizing fitness differences (-), i.e.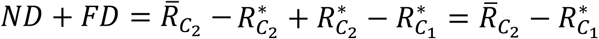.

## Appendix C: Fourier analysis of predator-prey oscillations

We found two different types of population dynamics in case of predator coexistence. For most parameter combinations, simple predator-prey oscillations emerged with the increase of the prey followed by an increase in the two predators (e.g. Figure 5E). For some other parameter combinations, the competition between *C*_1_ and *C*_2_ for the shared prey generated much more complex predator-prey dynamics (e.g. Figure 5F). To evaluate the differences in the population dynamics in more detail, we performed a discrete Fourier transformation by using the *fft* function implemented in MATLAB, version 7.13 (The MathWorks Inc., Natick, MA, 2011).

Our results show that for the simple predator-prey oscillations, most of the variation in the species’ abundances can be attributed to a single dominate time scale (Figure C1A, C and E). Here, the two predators are strongly synchronized, exhibiting a negligible phase-difference of 0.03 π (cf. Figure C1B, D). In contrast, for the more complex population dynamics, our analysis reveals two dominant time scales (Figure C2A, C and E), which can be most clearly seen from the amplitude spectrum of *C*_1_ (Figure C2C). At the low frequency, the population dynamics of the two predators exhibit a phase difference of 0.74 π (cf. Figure C2B, D). Hence, their abundances cycle almost anti-synchronously, meaning that an increase in the abundance of *C*_1_ coincides with a decrease in the abundance of *C*_2_ and *vice versa* (Figure C2D). In contrast to the prey (*R*) and *C*_1_, *C*_2_ showed hardly any variation in its abundance at the high frequency (Figure C2F), although this shorter time scale in the population dynamics is generated by the predator-prey interaction between *R* and *C*_2_. This discrepancy can be explained by the fact that the destabilization of the predator-prey oscillations by *C*_2_ selects immediately for the recovery of *C*_1_ (Figure C2B).

**Figure C1:**
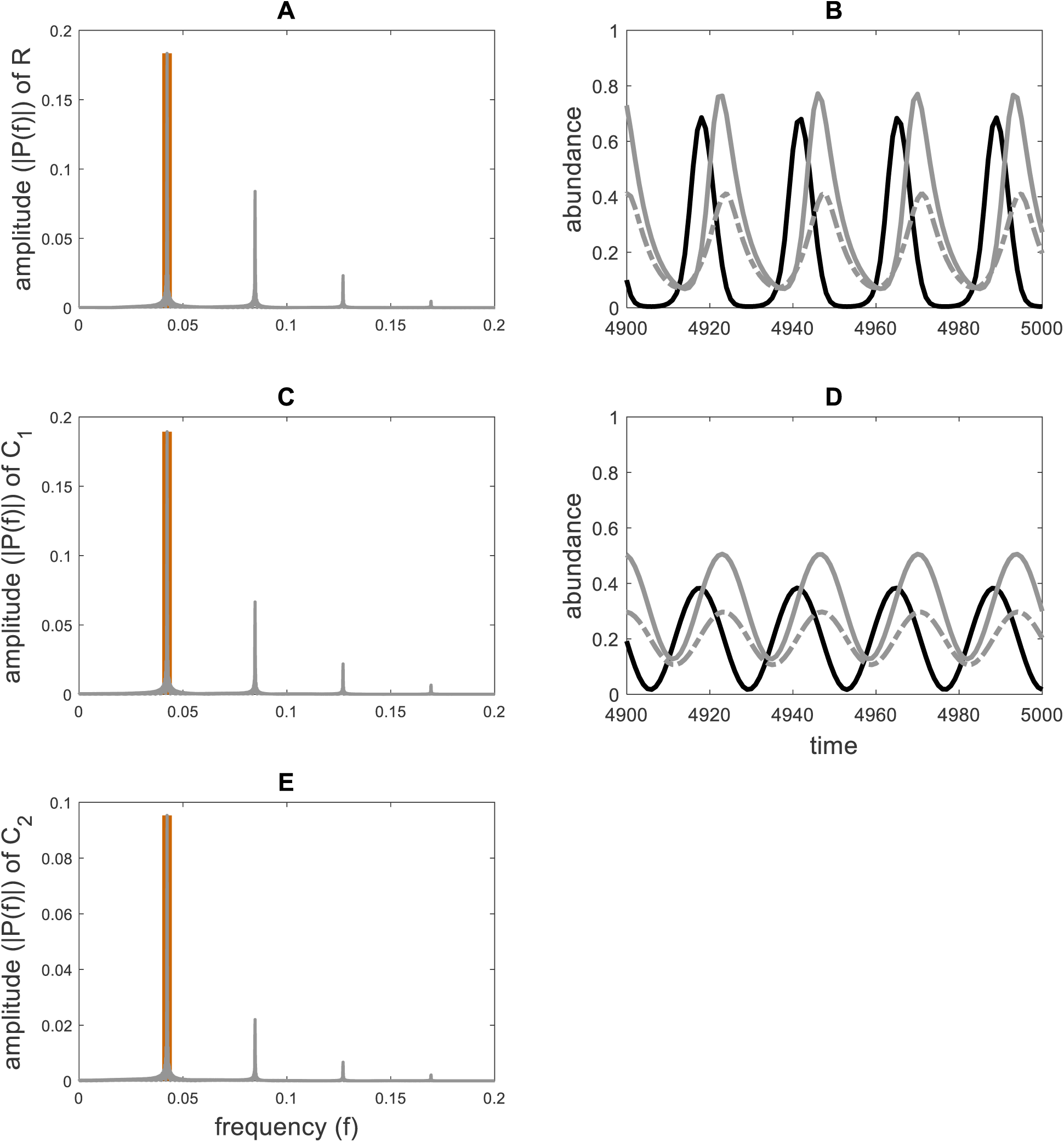
Frequency Domain and Time Domain of the population dynamics showing simple predator-prey oscillations (cf. Figure 4C). Amplitude spectrum (grey) of the time series of the prey (R), predator one (*C*_1_) and predator two (*C*_2_) are shown in panel A, C and E, respectively. The orange vertical line indicates the dominate time scale of the population dynamics. Panel B shows the full time series of R (black solid line), *C*_1_ (grey solid line) and *C*_2_ (grey dashed-dotted line). The population dynamics at the dominant time scale are plotted in panel D based on the calculated discrete Fourier transform.

**Figure C2:**
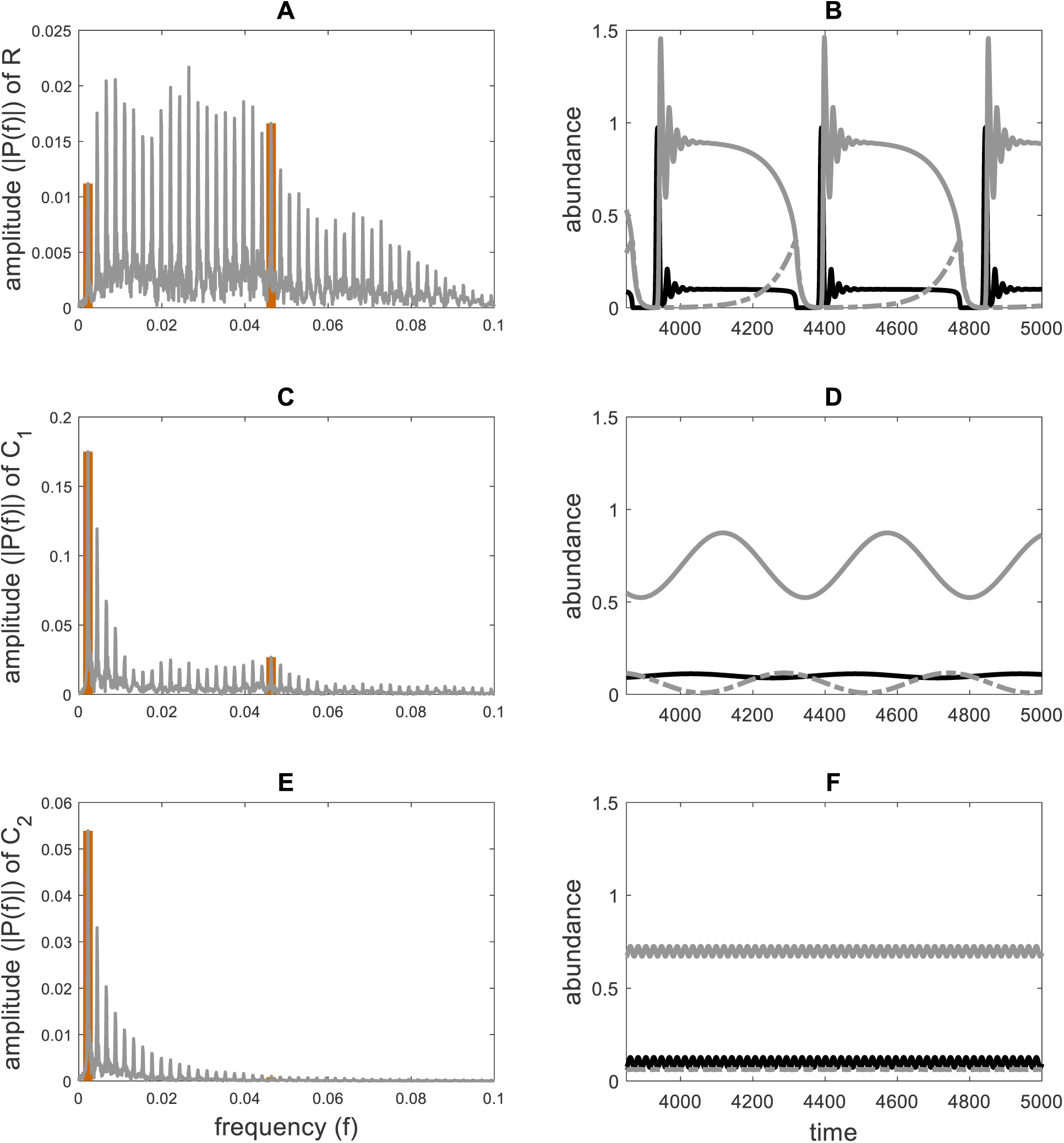
Frequency Domain and Time Domain of the population dynamics showing complex predator-prey oscillations (cf. Figure 4D). Amplitude spectrum (grey) of the time series of the prey (R), predator one (*C*_1_) and predator two (*C*_2_) are shown in panel A, C and E, respectively. The orange vertical lines indicate the two dominate time scales of the population dynamics. Panel B shows the full time series of R (black solid line), *C*_1_ (grey solid line) and *C*_2_ (grey dashed-dotted line). The population dynamics at the longer and shorter time scales are plotted in panel D and F, respectively, based on the calculated discrete Fourier transform.

## Appendix D The opposing effects of nutrient retention and nutrient recycling on the stability of predator-prey dynamics

Our results presented in the main text and Appendix A demonstrate that the retention of nutrients in the predator biomass stabilizes predator-prey dynamics because it effectively reduces the carrying capacity of the prey. In contrast, previous studies have shown that the recycling of nutrients from dead organisms destabilizes predator-prey dynamics (e.g. DeAngelis 1992; Kooi et al. 2002). Hence, non-consumptive effects on the prey may enhance or reduce the stability of food webs, depending on whether the stabilizing effects of nutrient retention or the destabilizing effects of nutrient recycling prevail.

To clarify the relative importance of nutrient retention and nutrient recycling for system stability we extended the predator-prey model used in the main text by two detrital components, thereby incorporating a time-lag in the recycling of nutrients from dead predators and prey (cf. DeAngelis 1992). We track the recycling from dead predators and prey separately because they generally contain different amounts of nutrients per unit of biomass. For convenience, we consider here one predator species only. Hence, the amount of dissolved inorganic nutrients (*N*) and the biomasses of the dead predators (*D*_*C*_) and prey (*D*_*R*_) and the living predators (*C*) and prey (*R*) are changing over time according to the following set of equations:

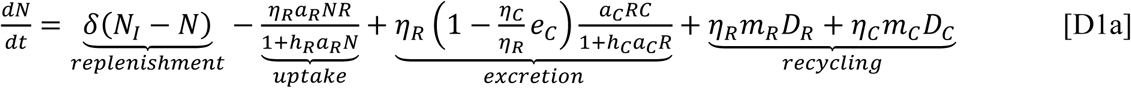

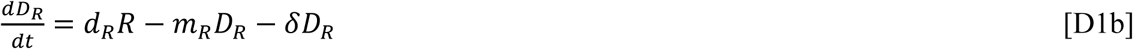

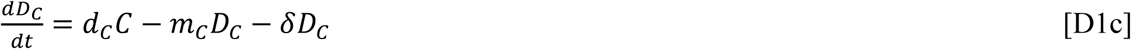

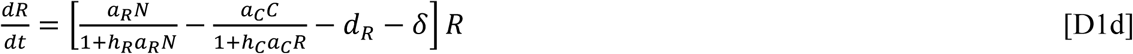

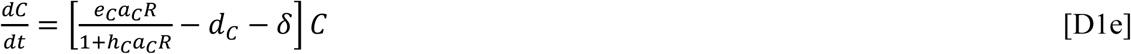

In contrast to eq. [1] of the main text, nutrient recycling does not occur immediately but rather passes through a detrital component. The time needed for the recycling of nutrients contained in the dead organisms into the inorganic nutrient pool scales with 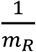 and 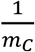 for the prey and predators, respectively. A complete description of the other terms and parameters can be found in the methods.

The predator-prey system described by eq. [D1] is mass balanced. Hence, the total amount of nutrients is given by *N*_*T*_ = *N* + *η*_*R*_(*R* + *D*_*R*_) + *η*_*C*_(*C* + *D*_*C*_). As a result, we can follow the general approach described in the methods and reduce the number of state variables of eq. [D1]. The dimensionally reduced predator-prey system is then given as follows:

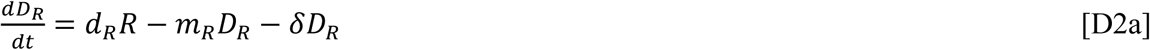

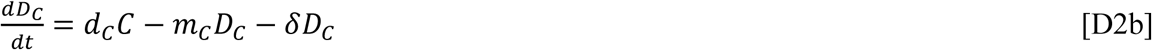

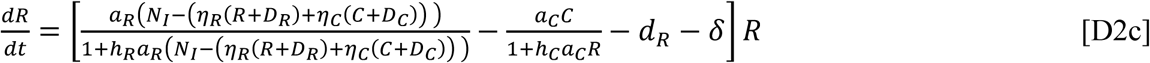

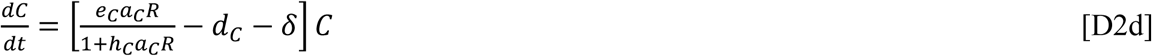

To further simply the analysis of our predator-prey system described by eq. [D2] we now consider a separation of time scales between the predator-prey dynamics (slow dynamics) and the recycling of nutrients from dead organisms (fast dynamics) (cf. Kooi et al. 1998b). Hence, we set 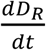 and 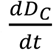 always equal to zero, and then substitute the resulting algebraic expression into eq. [D2c] (cf. O’Dwyer 2018). This results in the following approximation of our predator-prey system:

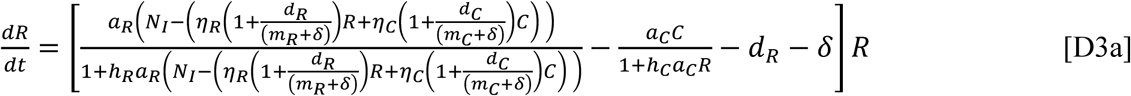

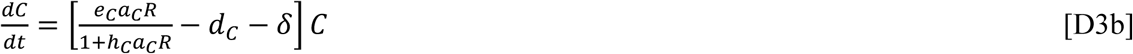

In the main text and Appendix A we showed that an increase in *η*_*R*_ and *η*_*C*_ has a stabilizing effect on the predator-prey dynamics. Consequently, an increase in *m*_*R*_ and *m*_*C*_ will have a destabilizing effect on the population dynamics of our predator-prey system described by eq. [D3] because *m*_*R*_ and *m*_*C*_ scale inversely with *η*_*R*_ and *η*_*C*_, respectively. Hence, our results confirm previous findings by e.g. DeAngelis (1992) and Kooi et al. (2002) that nutrient recycling has a destabilizing effect on the predator-prey dynamics (Figure D1). This happens because nutrient recycling effectively enhances the carrying capacity of the prey (cf. eq. D3a). In fact, in the absence of nutrient recycling (endogenously driven nutrient replenishment), i.e. *m*_*R*_ = 0 and *m*_*C*_ = 0, and dilution (exogenously driven nutrient replenishment), i.e. *δ* = 0, the amount of nutrients in the living predator and prey biomasses will decrease over time because the nutrients accumulate in the detrital compartments.

**Figure D1:**
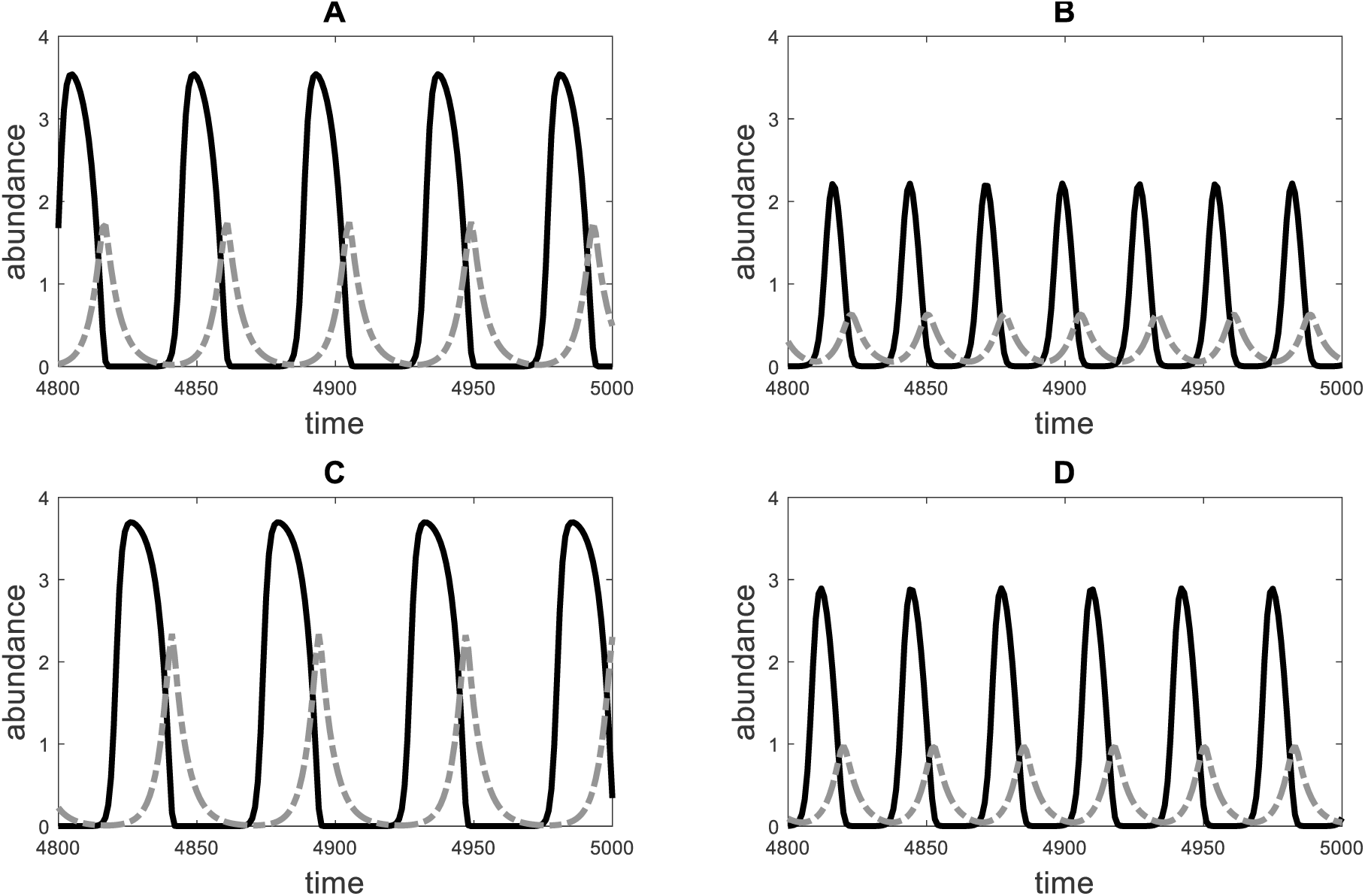
The predator (grey dashed-dotted line) and prey (black solid line) dynamics of the model defined by eq. [D3] in dependence of the rate of nutrient recycling (m_C_) and the nutrient to carbon ratio of the predators (η_C_). The parameter m_C_ equals 0.1 in panels A) and C) and 0.8 in panels B) and D) whereas the parameter η_C_ equals 0.25 in panels A) and B) and 1 in panels C) and D). Other parameter values are: d_R_ = 0; d_C_ = 0.15; a_R_ = 1; a_C_ = 2; h_R_ = 0; h_C_ = 2; e_C_ = 1; η_R_ = 0.25; δ = 0.05; N_I_ = 1.

